# Integrated analysis of Xist upregulation and gene silencing at the onset of random X-chromosome inactivation at high temporal and allelic resolution

**DOI:** 10.1101/2020.07.20.211573

**Authors:** Guido Pacini, Ilona Dunkel, Norbert Mages, Verena Mutzel, Bernd Timmermann, Annalisa Marsico, Edda G Schulz

## Abstract

To ensure dosage compensation between the sexes, one randomly chosen X chromosome is silenced in each female cell in the process of X-chromosome inactivation (XCI). XCI is initiated during early development through upregulation of the long non-coding RNA Xist, which mediates chromosome-wide gene silencing. Cell differentiation, Xist upregulation and silencing are thought to be coupled at multiple levels to ensure inactivation of exactly one out of two X chromosomes. Here we perform an integrated analysis of all three processes through allele-specific single-cell RNA-sequencing. Specifically, we assess the onset of random XCI with high temporal resolution in differentiating mouse embryonic stem cells, and develop dedicated analysis approaches. By exploiting the inter-cellular heterogeneity of XCI onset, we identify Nanog downregulation as its main trigger and discover additional putative Xist regulators. Moreover, we confirm several predictions of the stochastic model of XCI where monoallelic silencing is thought to be ensured through negative feedback regulation. Finally, we show that genetic variation modulates the XCI process at multiple levels, providing a potential explanation for the long-known Xce effect, which leads to preferential inactivation of a specific X chromosome in inter-strain crosses. We thus draw a detailed picture of the different levels of regulation that govern the initiation of XCI. The experimental and computational strategies we have developed here will allow us to profile random XCI in more physiological contexts, including primary human cells in vivo.

## Introduction

Female mammals carry two X chromosomes, while males have an X and a Y. Gene dosage differences between the sexes for X-linked genes are mostly compensated through X-chromosome inactivation (XCI). In this process, each female cell will silence one randomly chosen X chromosome in a cell-autonomous fashion (Galupa and Heard, 2015). Since XCI induces differential gene activity at two genetically identical chromosomes in the same nucleus, it is an important model for epigenetic gene regulation. The regulatory principles that allow the two X chromosomes to assume opposing activity states are only starting to be elucidated (Mutzel and Schulz, 2020).

A subset of X-linked genes are incompletely silenced on the inactive X chromosome and can thus escape XCI (Balaton and Brown, 2016). These escape genes are thought to contribute to phenotypic differences between the sexes, including susceptibility to various pathologies, such as autoimmune diseases (Snell and Turner, 2018). Moreover, inter-individual variability, for example with respect to the severity of X-linked diseases, is observed in cases where the choice of the inactive X chromosome is skewed through genetic polymorphisms (Peeters et al., 2016). Whether and to what extent differences in silencing efficiency and escape propensity driven by genetic variation might also contribute to phenotypic variability in humans remains unknown.

XCI is established during early embryonic development in a complex multi-step process. It is initiated by upregulation of the long non-coding RNA Xist, the master regulator of XCI (Galupa and Heard, 2015). Xist will coat the X chromosome, from which it is expressed, and will initiate chromosome-wide gene silencing by recruiting a series of silencing complexes, ultimately leading to heterochromatinization of the entire chromosome (Brockdorff et al., 2020). To ensure inactivation of a single X chromosome, Xist expression must be restricted to exactly one out of two alleles, in a monoallelic and female-specific fashion. While the majority of cells directly upregulate Xist monoallelically, we and others have recently shown that a subset of cells transiently express Xist from both chromosomes in a biallelic manner, which is subsequently converted to a monoallelic state (Mutzel et al., 2019; Sousa et al., 2018). The current model is thus that Xist is initially upregulated independently on each chromosome in a stochastic fashion and that establishment of a monoallelic state is then ensured through negative feedback regulation (Mutzel and Schulz, 2020). This feedback is thought to be mediated by silencing of an essential X-linked Xist activator (Lyon, 1971; Monkhorst et al., 2008). It remains unknown however, to what extent transient biallelic Xist upregulation indeed induces gene silencing, which is a prerequisite for the proposed negative feedback to work. A prediction from this “stochastic model of XCI” is that accelerated upregulation of Xist from one allele for instance caused by genetic variation will lead to preferential inactivation of that chromosome.

Random XCI is initiated during early embryonic development around the time of implantation into the uterus, when cells exit the pluripotent state. A series of factors have been implicated in triggering developmental Xist upregulation, such as the Xist activator Rnf12, Xist’s repressive antisense transcript Tsix and a series of pluripotency factors, such as Nanog, Oct4, Sox2, Klf4 and Rex1/Zfp42 (Donohoe et al., 2009; Gontan et al., 2012; Jonkers et al., 2009; Lee and Lu, 1999; Navarro et al., 2008, 2010). Pluripotency factors are indeed downregulated concomitantly with Xist upregulation and loss-of-function perturbations have been shown to increase Xist expression for several of them (Donohoe et al., 2009; Gontan et al., 2018; Navarro et al., 2008). It remains unknown, however, which factors trigger Xist upregulation in the endogenous context and whether this is mediated through modulating the activity of other Xist regulators such as Rnf12 or Tsix.

The different processes governing XCI have been studied extensively and we slowly see a picture emerging of how inactivation of one randomly chosen X chromosome might be achieved. However, many studies have been performed in engineered systems to be able to investigate a specific step in isolation (Barros de Andrade E Sousa et al., 2019; Marks et al., 2015; Żylicz et al., 2019). As a consequence a series of questions remain unanswered, which must be investigated in the endogenous context of random XCI with sufficient cellular, allelic and temporal resolution. Those questions include (1) to what extent gene silencing occurs upon biallelic Xist upregulation, (2) how differences in speed of Xist upregulation will affect the choice of the inactive X and (3) which factors actually trigger Xist expression. We therefore set out to perform an integrated analysis of Xist upregulation and gene silencing with single-cell resolution, in a context where cells make a random choice between their two X chromosomes. To this end we profiled XCI with high temporal resolution using allele-resolved single-cell RNA-sequencing (scRNA-seq). We used differentiating mouse embryonic stem cells (mESC), the classic tissue culture model of XCI. Since the cell line used has been derived from a hybrid cross between distantly related mouse strains, we could distinguish the two X chromosomes through single nucleotide polymorphisms (SNPs) and in addition investigate how genetic variation affected the different steps that govern XCI.

Building on this high-quality dataset, we have developed an analytical framework to study the heterogeneity of Xist expression, XCI dynamics and differentiation over time at the single cell level. Specifically, we analyzed the full transcriptome heterogeneity throughout the estimated pseudotime and used RNA velocity to annotate lineage trajectories corresponding to the choice of the inactive X chromosome. By exploiting the variability in gene expression across cells we were able to recover known regulators of Xist, but could also identify potential novel Xist repressors and activators. We found that XCI is initiated on both chromosomes in a subset of cells through biallelic Xist upregulation, but is rapidly reversed to a monoallelic state, when silencing is initiated. By computing Xist expression and gene-silencing dynamics in an allele-specific manner, we discovered that genetic variation modulated XCI at multiple levels, including expression frequency and level of Xist as well as chromosome-wide silencing efficiency and escape propensity of individual genes. Finally, we validated these findings through an orthogonal experimental approach. Our study thus provides a detailed view on how random X inactivation is first established. The approaches we have developed will also be instrumental to profile endogenous XCI in other contexts, such as primary human tissues.

## Results

### X inactivation and X upregulation in differentiating mouse embryonic stem cells

At the onset of XCI, the inactive X chromosome is chosen in a random cell-autonomous process, resulting in a mixture of cells that have silenced the paternal or the maternal X chromosome, respectively. To investigate the onset of random XCI with single-cell resolution, we profiled female mESCs at multiple time points during differentiation by 2i/Lif withdrawal using scRNA-seq (Fig. 1A). To allow allele-specific transcript quantification, we used a F1 hybrid mESC line (TX1072), which has been derived by crossing two distantly related mouse strains, *Mus musculus domesticus* (C57BL6/J) and *Mus musculus castaneus* (Cast/EiJ), herein referred to as B6 and Cast, respectively (Schulz et al., 2014). At multiple time points during differentiation (0,1,2,3,4 days) we captured in total 1945 individual cells on a C1 microfluidics system (Fluidigm). Single-cell transcriptomes were profiled using the C1-HT protocol, which performs 3’-end counting using unique molecular identifiers (UMI) and thus enables strand-specific analyses. Around 0.4 Mio reads were sequenced per cell, resulting in a median number of 0.12 Mio unique UMI counts, which represent the detected mRNA molecules, covering a median number of 6090 genes (248 X-linked) per cell (Fig. S1A+B). After removing low-quality cells and cells that had lost one X chromosome, between 257-341 cells were retained per time point for further analysis (Fig. S1B-C, Supplemental Table S1).

**Figure 1.**
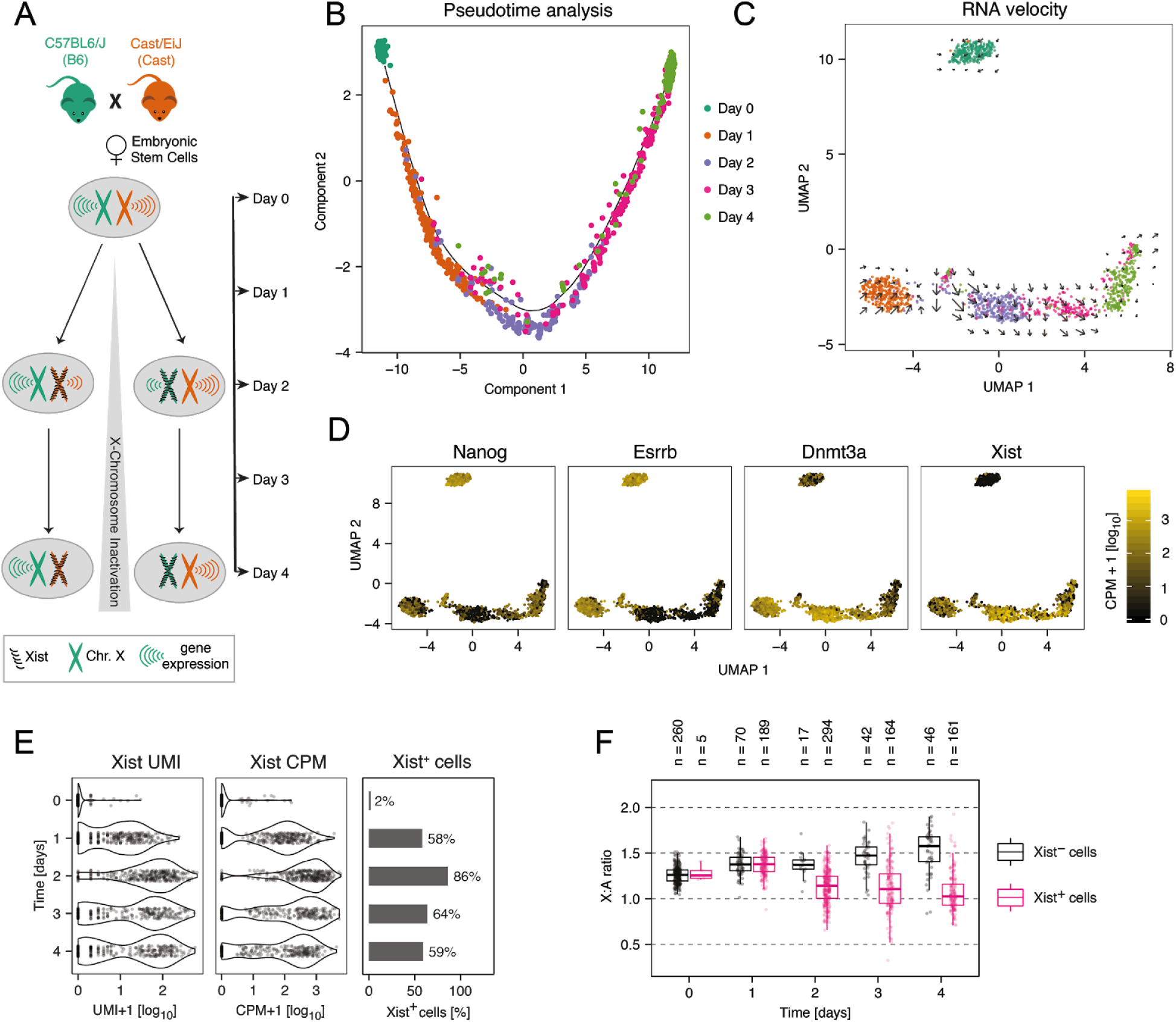
Profiling the onset of random XCI by scRNA-seq. **(A)** Schematic representation of the experimental setup. A female mESC line derived from the cross between a B6 (green) and a Cast (orange) mouse was differentiated for 4 days by 2i/Lif withdrawal and up to 400 single-cell transcriptomes were collected per time point. During the time course, cells initiate random XCI by monoallelic upregulation of Xist (black) from one randomly chosen allele, which will induce chromosome-wide gene silencing. **(B-C)** Pseudotime analysis (B) and UMAP embedding (C) based on the 500 most variable genes, with individual cells colored by measurement time. The black line in B represents the principal graph describing the pseudotime trajectory of the projected cells as computed by Monocole2 DDRTree method. Arrows in C indicate the predicted transcriptome change estimated through RNA velocity analysis. **(D)** UMAP embedding as in C with cells colored according to marker gene expression. **(E)** Distribution of Xist expression across cells, either shown as the number of UMI counts (=number of molecules, left), the normalized CPM value (middle) or the percentage of Xist-positive (>5 Xist UMI counts) cells (right). **(F)** Boxplot of the X-to-autosome expression ratio in Xist-positive (pink) and Xist-negative cells (black). The central mark indicates the median, and the bottom and top edges of the box indicate the first and third quartiles, respectively. The top and bottom whiskers extend the boxes to a maximum of 1.5 times the interquartile range. Dots represent individual cells, the number of cells in each group is given on top.

In a first step we analyzed how the cells’ transcriptomes changed over the time course measured. Dimensionality reduction using UMAP (McInnes et al., 2018) and pseudotime analysis with Monocle (Trapnell et al., 2014) revealed that undifferentiated mESCs clustered distantly from their differentiating derivatives, suggesting that the change of culture condition induced a major change in their transcriptomes (Fig. 1B+C). Pseudotime analysis resolved the sampling time of most cells during differentiation (Fig. 1B). It also highlighted differentiation heterogeneity between cells, which became maximal at day 2-3 with a subset of cells clustering close to 1 day-differentiated cells, suggesting a differentiation block (Fig. 1B+C). To assess the lineage trajectory of each cell we performed an RNA velocity analysis, which estimates transcriptional activity from reads aligning to introns and uses that information to predict how mature mRNA expression will change in the future (Fig. 1C, arrows, Fig. S1D) (La Manno et al., 2018). The results suggested that cells moved along a single differentiation trajectory. This was confirmed by an analysis of marker genes revealing that naive pluripotency factors such as Nanog and Esrrb were downregulated while Dnmt3a, a marker of primed pluripotency was upregulated (Fig. 1D).

Analysis of Xist expression in the UMAP projection showed the expected upregulation during differentiation, but also revealed marked heterogeneity with a subset of cells expressing no or low levels of Xist (Fig. 1D, right). Xist was detected with >5 UMI counts (Xist-positive cells) in only 2% of undifferentiated mESCs, but in 58-86% of cells during differentiation, with the expression level varying strongly between cells and between time points from around 10 molecules at day 1 to >100 molecules at later stages (Fig. 1E). Interestingly, Xist expression appeared to be maximal at day 2 and decreased at later time points. We estimated the detection rate in our data set to lie around 30%, given that the number of mRNAs present in an ES cell has been estimated to be around ∼400,000 molecules (Carter et al., 2005), ∼120,000 of which we detected per cell (Fig. S1A+B). The actual mean copy number of the Xist RNA in Xist-expressing cells would thus increase from 79 at day 1 to 243-314 at the later time points, which is in good agreement with a previous estimate of ∼300 molecules(Sun et al., 2006).

When analyzing the read distribution along genes, we noticed an unusual read pattern for Xist. Instead of the expected 3’ bias (see examples in Fig. S2A-C), we observed a strong peak in Xist’s first exon, but did not detect a robust signal at its 3’-end (Fig. S2D-E). A similar read pattern was also found in a previously published data set, which used CEL-seq2, a different 3’-end scRNA-seq method, to profile mouse fibroblasts (Fig. S2F)(Hashimshony et al., 2016). Inspection of the sequence upstream of the peak revealed a genomically encoded poly-adenine (polyA) stretch within the Xist RNA, which likely served as a template to prime reverse transcription at this position (Fig. S2D). The fact that read density in Xist’s first exon showed the expected temporal profile (upregulation over time, Fig. 1E) suggested that Xist was nevertheless correctly quantified in our data set, even though its polyA tail appeared to be inaccessible to the reverse transcription reaction.

To assess whether we could observe Xist-induced gene silencing, we estimated expression from the X chromosome relative to autosomal genes (X:A ratio, Fig. 1F). We classified cells as Xist-positive, if >5 UMI counts were assigned to the *Xist* gene and as Xist-negative, if Xist was not detected in the cell. Starting from day 2 of differentiation we observed a clear downregulation of X-linked genes in Xist-positive cells, from a median X:A ratio of 1.25 across all cells before differentiation to 1.03 at day 4 in Xist-positive cells (Fig. 1F, p<2.2*10^−16^, Mann-Whitney U two-sided test). Interestingly, the X:A ratio increased to 1.58 over time in Xist-negative cells, which presumably failed to initiate XCI (p<6.2*10^−14^, Mann-Whitney U two-sided test). Such global upregulation of X-linked genes has been reported previously in differentiating male mESCs and in male pre- and post-implantation embryos in vivo (Borensztein et al., 2017a; Larsson et al., 2019; Lin et al., 2007; Mohammed et al., 2017; Wang et al., 2016). This process termed X upregulation is thought to have evolved to compensate for the loss of Y-chromosomal genes (Ohno’s hypothesis), but is still controversial (Disteche, 2016).

### Allele-specific analysis of Xist expression patterns

To quantify Xist in an allele-specific manner, we mapped the sequencing results to the mouse reference genome with masking all SNPs present in the TX1072 cell line, and counted the reads that could be assigned to one or the other allele. Around 4% of all reads could be mapped in an allele-specific manner (Fig. S1A). Allele-specific analysis of Xist expression revealed the expected monoallelic pattern after 3-4 days of differentiation, where the vast majority of cells expressed Xist either from the B6 or from the Cast chromosome (Fig. 2A, orange+green). At day 1 and 2 of differentiation by contrast, a subpopulation expressed Xist from both alleles (Fig. 2A+B, pink). To quantify the Xist expression patterns, we classified all cells as follows (Supplemental Table S2). Cells with ≤ 5 allele-specific Xist UMI counts were termed Xist-low (Fig. 2A+B, grey) and those where Xist was not detected at all Xist-negative (black). Cells with >5 Xist UMI counts were classified as monoallelic, if all counts mapped to the same allele (MA-B6 or MA-Cast, orange, green, Fig. 2A+B). If more than 80% of Xist counts were assigned to one allele, a cell was labelled as skewed (Fig. 2A+B, light pink) and if at least 20% were assigned to each allele it was defined as biallelic (BA, dark pink).

**Figure 2.**
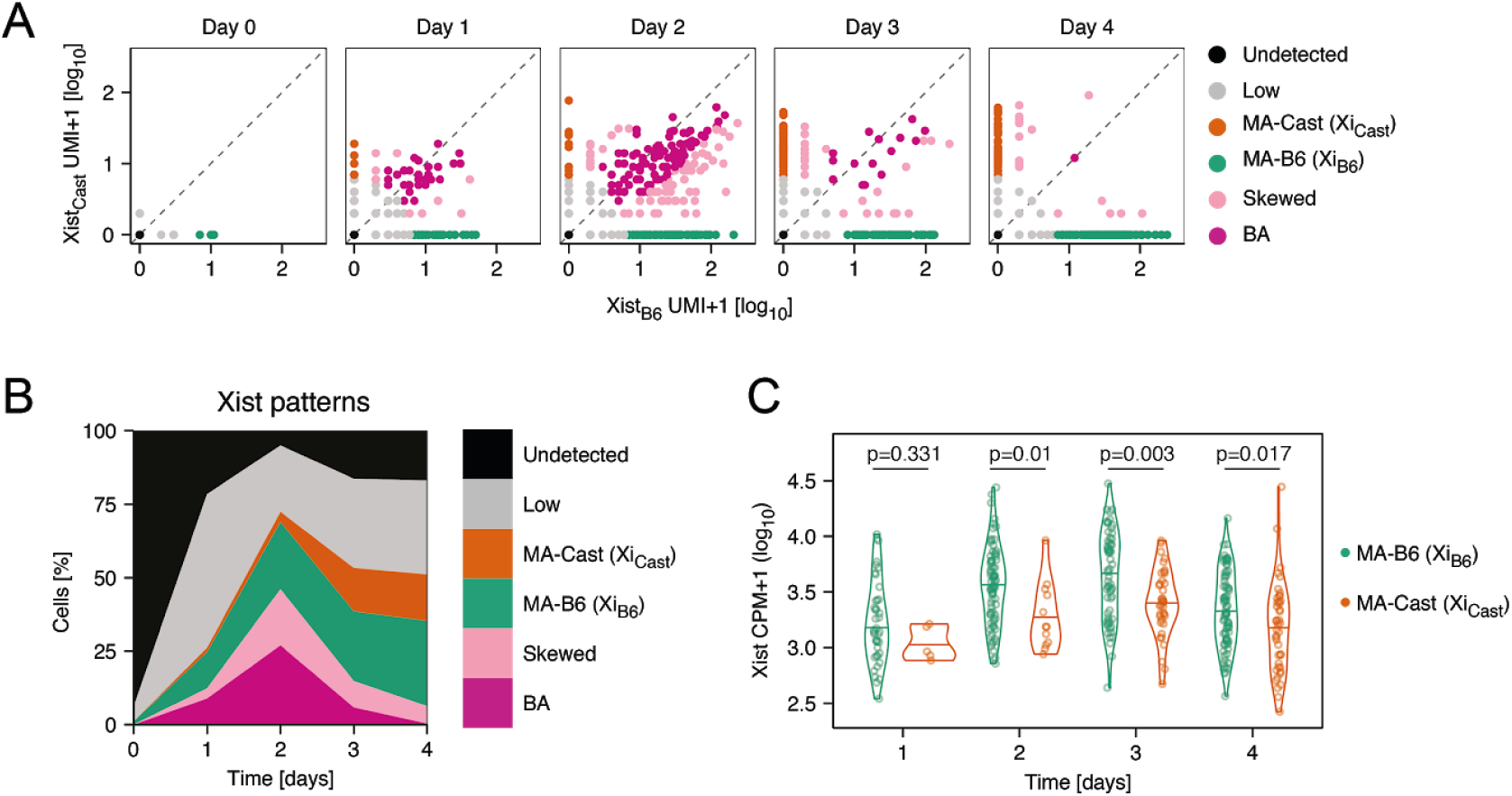
Allele-specific analysis of Xist expression. **(A)** Scatter plot showing Xist UMI counts mapping to the B6 and Cast chromosome in individual cells. Cell coloring indicates the Xist pattern classification, where Xist detection from only one allele is termed monoallelic (MA, orange, green). Detection of both chromosomes is termed biallelic (BA, pink), if at least 20% of UMI counts come from each allele, or termed skewed (light pink) if one allele contributes <20% of reads. Cells with ≤5 allele-specific UMI counts were defined as Xist-low (grey) and cells without Xist detection are shown in black. **(B)** Relative distribution of Xist patterns as in (A) across time. **(C)** Violin plot comparing Xist expression levels from the B6 (green) and Cast chromosomes (orange) in Xist-MA cells. The horizontal line indicates the median value and p-values of a Mann-Whitney U two-sided test are shown.

At day 1 and 2 at least half of Xist-positive cells (>5 UMI counts) expressed both alleles (Fig. 2B, light and dark pink), while at later time points the majority exhibited a monoallelic expression pattern (Fig. 2B, green, orange). We and others have recently found such transient biallelic Xist upregulation at the onset of random XCI in vivo (Mutzel et al., 2019; Sousa et al., 2018). Overall, more cells upregulated Xist from the B6 than from the Cast allele (29% vs 16% at day 4), which is in agreement with the previously reported preferential inactivation of the B6 allele in B6xCast F1 hybrid cells due to differing X-controlling elements (Xce) (Chadwick et al., 2006; Plenge et al., 2000). Interestingly, the B6 allele appeared to upregulate Xist faster and reached higher levels than the Cast allele (median UMI 31 vs 16 at day 4) (Fig. 2B+C). This observation is in agreement with the prediction of the stochastic model of XCI onset that faster Xist upregulation from one allele will result in preferential X inactivation of that chromosome and could account for the Xce effect in this hybrid context (Monkhorst et al., 2008; Mutzel and Schulz, 2020). In summary, our allele-specific analysis of Xist expression revealed initial biallelic upregulation, which was then resolved to a monoallelic state with preferential inactivation of the B6 chromosome.

### Identification of putative Xist regulators

At all time points we observed marked heterogeneity across cells with regard to Xist expression, which varied over more than 2 orders of magnitude with no Xist being detected in a subset of cells (Fig. 1E). We reasoned that we could make use of the variability within the generated single-cell transcriptome data to address the question of how Xist upregulation is triggered at the onset of XCI. We would expect that the responsible Xist activators or repressors should correlate positively or negatively with Xist across cells. We used two different approaches to identify such candidate regulators, based on differential expression and correlation analysis, respectively, to ensure robustness of the results (Supplemental Table S3). For the differential expression analysis we compared at each time point cells expressing low and high levels of Xist. To classify cells according to Xist expression, we performed k-means clustering to identify 7 clusters and defined cells in the 3 clusters with the highest expression as Xist-high, and cells in the 3 lowest clusters as Xist-low (Fig. 3A, for details see computational methods). Cells in the intermediate cluster were excluded from the analysis. We then identified differentially expressed genes (DEGs) between Xist-low and Xist-high cells at each time point using the MAST method (Finak et al., 2015). While only two DEGs (FDR<0.05) were detected at day 1, the number of both autosomal and X-linked DEGs increased over time up to 274 and 82, respectively, at day 4 (Fig. 3B, Fig. S3A). The vast majority of X-chromosomal DEGs were expressed at reduced levels in Xist-high cells due to Xist-mediated gene silencing, and their numbers steadily increased between day 2 and 4 from 22 to 81 (Fig. 3B, Fig. S3, right, blue). The opposite pattern was only found for Pim2, which encodes an oncogenic kinase that cooperates with the Myc transcription factor (Mondello et al., 2014).

**Figure 3.**
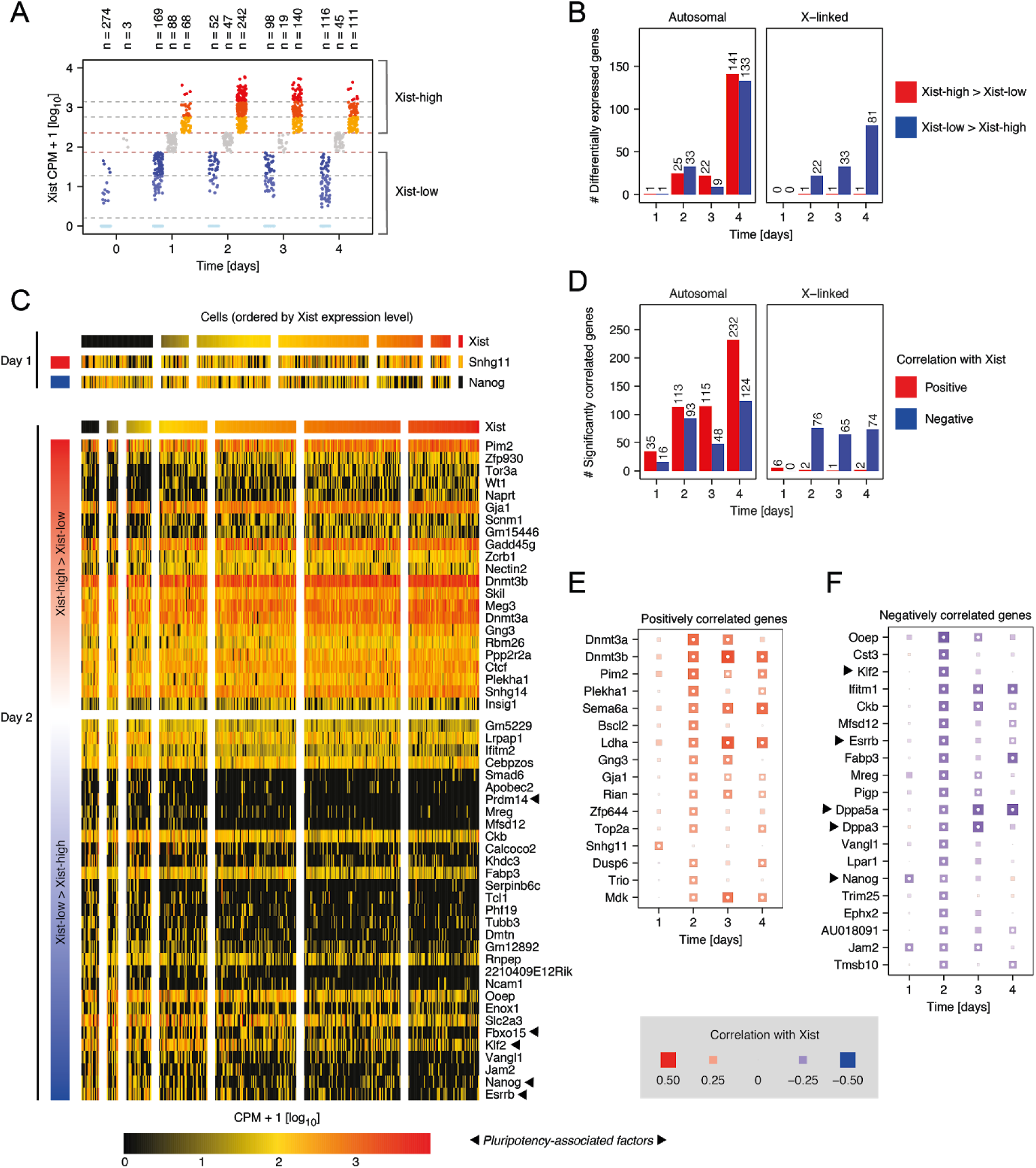
Identification of putative Xist regulators. **(A)** Cell classification according to Xist expression levels. The three highest and lowest cell clusters from a K-means (k=7) clustering were defined as Xist-high (yellow/red) and Xist-low (blue), respectively. **(B)** Number of differentially expressed genes (DEGs), excluding Xist, between Xist-high and Xist-low cells on autosomes (left) and on the X-chromosome (right) (BH correction, FDR<0.05). **(C)** Heatmaps showing expression of DEGs (rows) with absolute fold change between Xist-high and Xist-low cells above 1.5 at day 1 (top) or day 2 (bottom) in single cells (columns). Cells are ordered by Xist expression level and grouped according to the clustering shown in A. Genes are ordered by decreasing fold change. X-linked genes with Xist-low>Xist-high are not shown. **(D)** Number of genes, whose expression is positively (red) or negatively (blue) correlated with Xist expression (Spearman’s correlation test and BH correction, FDR<0.05) across cells of the same time point. **(E-F)** Spearman’s correlation coefficients with Xist for positively correlated genes (E) and negatively correlated autosomal genes (F). Only genes that exhibit a significant correlation (FDR<0.05) at day 1 or 2 are shown, with an absolute Spearman’s correlation coefficient >0.25. Size and color indicate the correlation coefficient as indicated. White dots represent significant correlation (Spearman correlation test and BH correction, FDR<0.05).

With XCI progression the number of autosomally encoded DEGs increased strongly up to 274 at day 4, presumably due to global X-dosage effects such as modulation of the differentiation-promoting MAPK signalling pathway and DNA hypomethylation (Schulz et al., 2014; Song et al., 2019; Zvetkova et al., 2005). At later time points, genes might thus be differentially expressed either because their expression is affected by XCI or because they regulate XCI. To specifically identify XCI regulators, we focused on day 1 and 2 of differentiation, when X-dosage effects were less pronounced because silencing was just being initiated (see below). On day 1, only 2 DEGs were detected, the known Xist regulator Nanog and the lncRNA Snhg11 (Fig. 3C, top). On day 2, 58 autosomal DEGs were found. Among the genes that were significantly higher expressed in Xist-low cells, were the pluripotency factors Nanog, Klf2 and Prdm14 (2.2, 1.9, 1.2 log2 fold change), which have previously been implicated in Xist repression (Gillich et al., 2012; Navarro et al., 2008; Payer et al., 2013). Moreover, several other pluripotency-associated factors such as Esrrb and Fbxo15 were identified as potential Xist repressors (2.9, 1.9 log2 fold change) (Martello and Smith, 2014; Okita et al., 2007). Among the putative activators we found several genes with potential roles in transcriptional regulation or signalling, such as the transcription factors Wt1 and Gm15446, DNA methyltransferases Dnmt3a and Dnmt3b, the splicing factor Zcrb1, and signalling factors Gadd45g and Skil, which modulate MAPK and TGFb/Smad signalling respectively (Ecco et al., 2016; Greenberg and Bourc’his, 2019; Jahchan and Luo, 2010; Salvador et al., 2013; Wagner et al., 2003).

In a second approach we calculated Spearman’s correlation coefficient between the normalized expression of Xist and all other detected genes across cells of each time point separately, an approach that has been successfully applied to scRNA-seq data previously (Kolodziejczyk et al., 2015). Also here the number of significantly correlated genes (FDR<0.05) increased over time and the correlation was mostly negative for X-chromosomal genes due to Xist-induced silencing (Fig. 3D). To identify relevant Xist activators (Fig. 3E) and Xist repressors (Fig. 3F) we focused on genes with an absolute correlation coefficient above 0.25 at day 1 or 2 and excluded negatively correlated X-linked genes from our analysis. At day 1, only Jam2, a stem cell marker (Sakaguchi et al., 2006), and Snhg11 and Nanog, which were also identified in the differential expression analysis (Fig. 3C), were significantly correlated with Xist. At day 2 the highest positive correlation was observed for Dnmt3a and Dnmt3b and the X-linked factor Pim2, all of which we had also identified as DEGs (Fig. 3C). In addition, the transcription factor Zfp644 and signalling genes Gng3, Dusp6 and Trio were identified as putative activators (Fig. 3E). Among the top negatively correlated genes we found again several pluripotency factors, such as Klf2, Esrrb, Nanog, Dppa5a and Dppa3 (Fig. 3F). Other potentially interesting factors were Vangl1, which modulates Wnt signalling and Trim25, an E3 ubiquitin ligase (Mentink et al., 2018; Munir, 2010). Interestingly, this analysis only identified a subset of the previously proposed regulators. A comprehensive analysis of genes that have been implicated in Xist regulation before, showed that Nanog, Klf2/4 and Prdm14 were correlated with Xist, while other pluripotency factors such as Oct4 and Sox2 were not (Suppl. Fig. S3B). The initial trigger of Xist upregulation thus seems to be Nanog, which was differentially expressed and negatively correlated with Xist already at day 1, with Klf2/4 and Prdm14 potentially contributing to a lesser degree. Another, newly identified putative Xist regulator potentially implicated in initial Xist upregulation is Snhg11. It might be involved in inducing Xist expression, since it was positively associated with Xist in both our approaches. This poorly studied lncRNA, which has been implicated in cancer warrants further investigation in the future (Huang et al., 2020).

### Global gene silencing dynamics

To integrate the analysis of Xist regulation with Xist-induced gene silencing, we next quantified global gene activity of the X chromosome in an allele-specific fashion. First, we calculated for each cell the fraction of X-chromosomal reads (excluding Xist) that mapped to the B6 allele (Fig. 4A). A B6 fraction around 0.5 before differentiation and in differentiating Xist-negative cells reflected equal activity of both X chromosomes. In Xist-expressing cells the distribution broadened over time until two distinct populations became visible at day 4, exhibiting B6 fractions approaching 0 or 1, respectively. This indicated that random X inactivation was initiated, where each cell silenced either the B6 or the Cast X chromosome. In the next step, we wanted to analyze the silencing dynamics in individual cells using the concept of RNA velocity (La Manno et al., 2018) with adaptation to the allele-specific analysis of XCI. To this end we quantified spliced and unspliced reads for all X-linked genes on each allele separately (Fig. 4B, Fig. S1D). The expected XCI trend was observed in both cases, with the Xist-expressing allele progressively reducing gene activity over time. This observation convinced us to model the allele-specific spliced and unspliced transcripts of each X-linked gene through the RNA velocity method (La Manno et al., 2018), to predict the future allelic B6 fraction of each cell. The predicted cell states were then projected onto the first two principal components of the B6 fractions for all X-linked genes (Fig. 4C). The first principle component (PCA 1) separated cells that silenced the Cast (orange) and the B6 chromosome (green) and RNA velocity revealed trajectories towards the silenced states.

**Figure 4.**
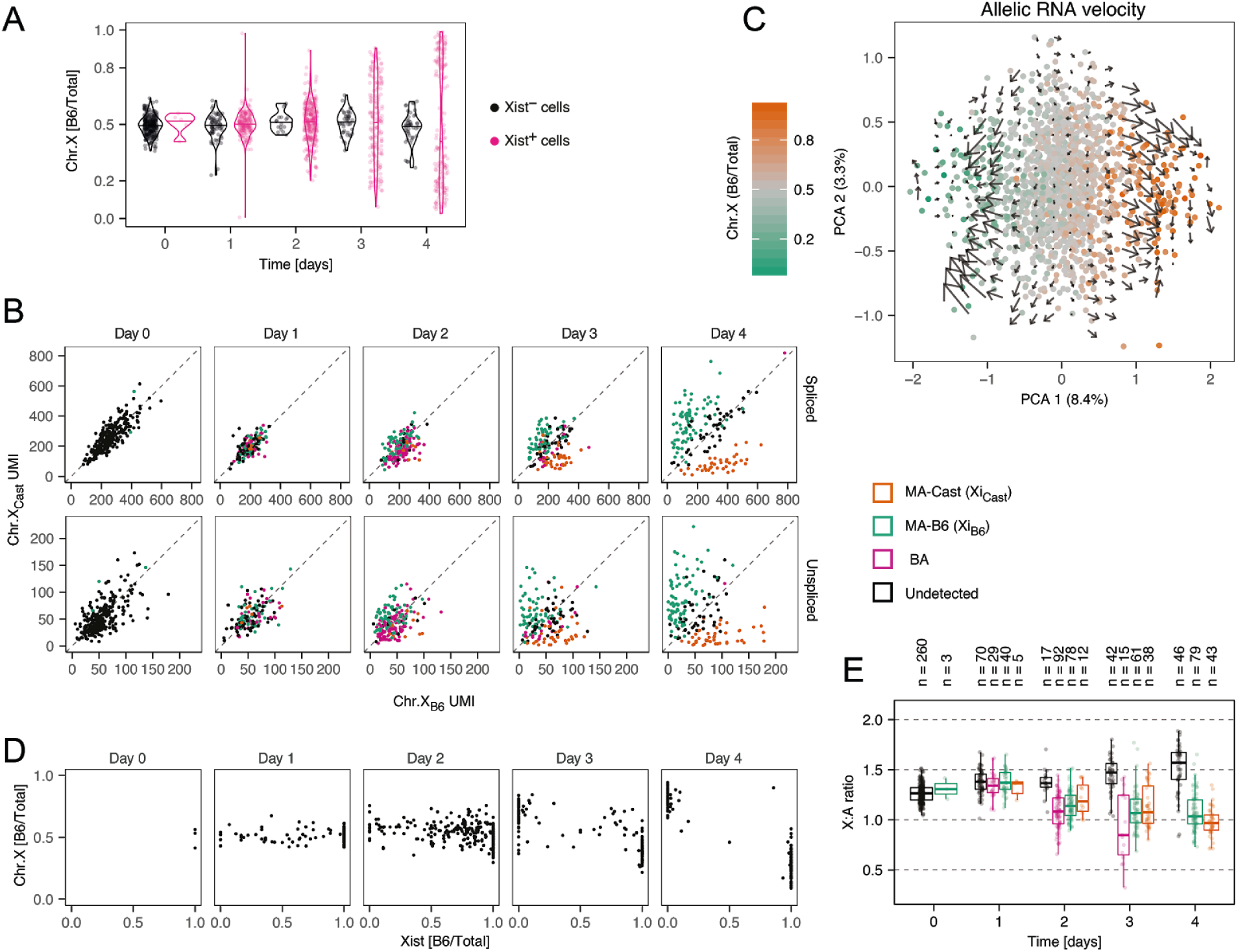
Chromosome-wide silencing dynamics. **(A)** Violin plot showing the distribution of the allelic expression ratio for the entire X chromosome excluding Xist, as the fraction of X-chromosomal reads mapping to the B6 chromosome, in Xist-positive (pink) and Xist-negative cells (black). Horizontal lines indicate the median values. **(B)** Scatter plots showing spliced (top) and unspliced (bottom) reads mapping to the X chromosome on the B6 and Cast alleles. Cells are colored by Xist expression pattern as in Fig. 2A. (**C)** The first two principal components, computed on the allelic expression ratio for all X-linked genes with the color indicating the chromosome-wide allelic ratio. Arrows indicate the predicted change of the X-linked allelic ratio based on RNA velocity analysis. **(D)** Scatter plot showing the X-chromosomal allelic ratio vs. Xist’s allelic ratio, for cells with >5 Xist allele-specific UMI counts. **(E)** Box plot showing the X:A expression ratio in cells assigned to different Xist classes based on allele-specific mapping as in Fig. 2A. Dots represent individual cells and cell numbers are indicated on top.

We next integrated the chromosome-wide silencing analysis with the allele-specific analysis of Xist expression, quantified as the fraction of Xist counts assigned to the B6 allele. We found that Xist expression from the B6 allele was associated with a skewing of X-chromosomal gene expression towards the Cast chromosome and vice versa at day 3 and 4 of differentiation, reflecting Xist-induced chromosome-wide gene silencing (Fig. 4D). Moreover, transient biallelic Xist expression was resolved to a monoallelic state at the same time when chromosome-wide silencing became visible (around day 3). This observation is in line with a putative role of biallelic silencing of an X-linked Xist activator in reversing biallelic Xist upregulation, which we have recently proposed (Mutzel et al., 2019). The extent of gene silencing in cells with biallelic Xist expression had however remained unknown. Since biallelic silencing cannot be investigated with the allele-specific approach used above, we instead assessed the X:A ratio in cells with different Xist expression patterns (Fig. 4E). At day 2 and 3 the onset of gene silencing was clearly visible in biallelic cells and silencing was even more pronounced than in cells with monoallelic Xist expression (p=0.01/0.08 at day 2/3, Mann-Whitney U two-sided test), suggesting that gene silencing was indeed initiated at both X chromosomes. In summary, allele-specific scRNA-seq can indeed be used to quantitatively assess the relationship between Xist expression and global gene silencing at the onset of random XCI with single cell resolution. In this way we could show that chromosome-wide gene silencing started around two days after Xist was initially upregulated and coincided with the biallelic-to-monoallelic transition for Xist.

### Gene- and allele-specific silencing dynamics

Since our time course experiment was performed in a highly polymorphic cell line, it also provided the opportunity to ask how genetic variation affected the efficiency or dynamics of gene silencing. When comparing XCI in MA-B6 and MA-Cast cells, we found that the Cast chromosome appeared to be silenced more efficiently (Fig. 5A), although we had found that Xist was upregulated more slowly and to lower levels at that chromosome (Fig. 2B+C). The fact that both Xist and other X-linked genes appeared to be preferentially detected from the B6 chromosome might suggest a technical artifact, such as a mapping bias towards the reference genome (B6). Although we detected a slight tendency for higher expression from the B6 chromosome (1493/1078 autosomal genes with higher expression on B6/Cast, 2288 genes unchanged, p=0.002, Fisher’s exact test), the fact that the median allelic ratio was 0.99-1.05 for autosomal genes and for X-linked genes in Xist-negative cells did not provide an indication for a strong mapping bias (Fig. S4). We thus concluded that genetic variation between the B6 and Cast alleles seemed to result in Xist being upregulated with higher probability and faster dynamics from the B6 X chromosome, while subsequent silencing was induced more rapidly on the Cast allele.

**Figure 5.**
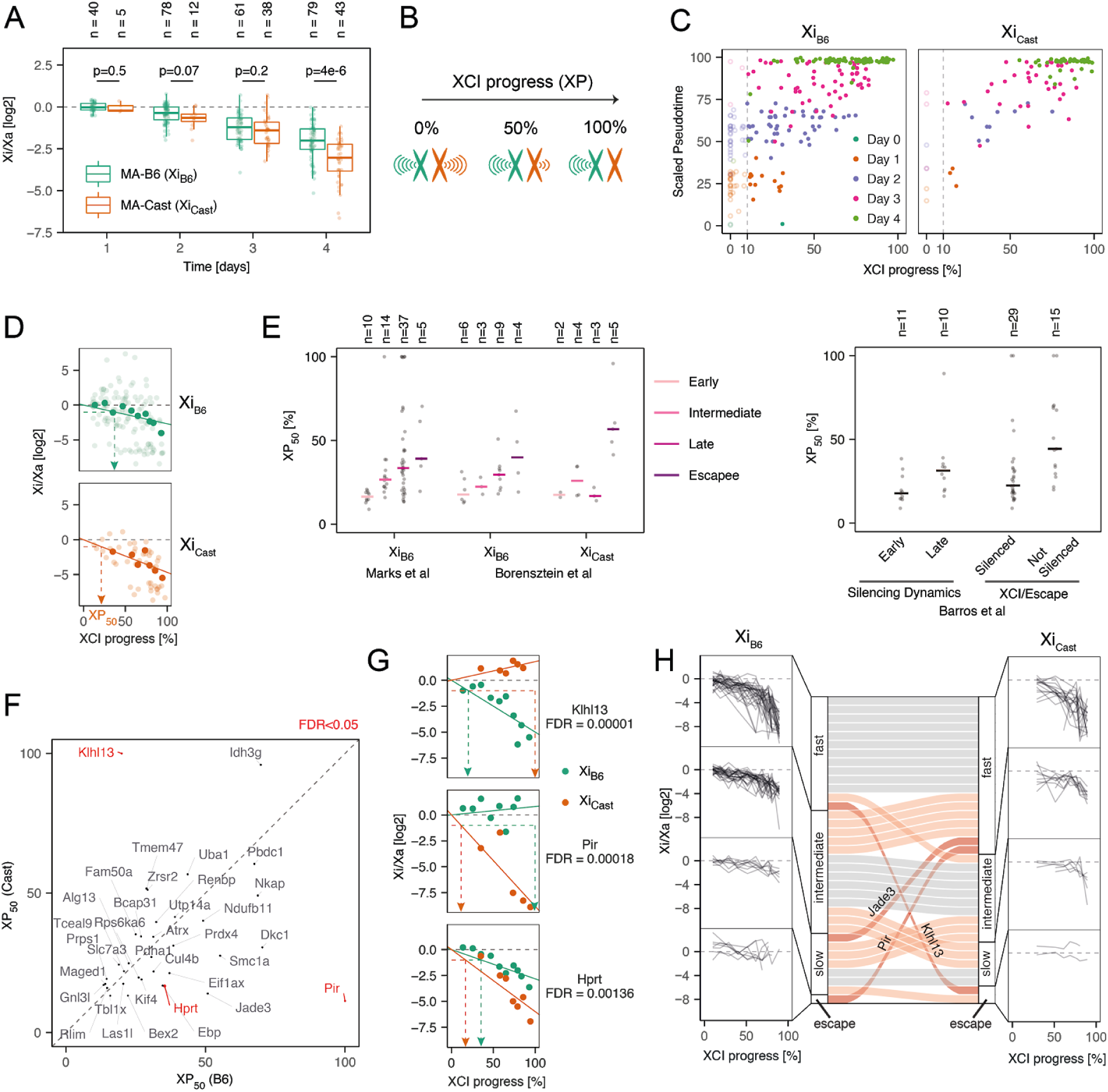
Comparison of allelic silencing dynamics. **(A)** Xi:Xa expression ratio for cells that inactivate the B6 (green) and Cast chromosome (orange), excluding reads mapping to Xist. Dots represent individual cells and cell numbers are indicated on top; p-values of a Mann-Whitney U two-sided test are shown. **(B)** Schematic representation of XCI progress (XP), defined as the percentage of silencing. **(C)** Comparison of XCI progress with scaled pseudotime as in Fig. 1B for MA-B6 (left) and MA-Cast cells (right). Cells are colored according to measurement time point. Only cells with XP>10% (vertical dashed line) were included in the differential silencing analysis in D-G. **(D)** Estimation of allele-specific silencing dynamics shown for an example gene (Eif1ax). Transparent dots indicate individual cells and solid dots the binned values for cells with similar XCI progress. The solid line shows the log-linear fit, used to estimate the XP_50_ value (dashed arrows), which describes the global XCI progress at which a given gene is silenced by 50%. **(E)** Comparison of the estimated XP_50_ values with previously determined silencing classes (Barros de Andrade E Sousa et al., 2019; Borensztein et al., 2017a; Marks et al., 2015). Dots represent individual genes and the horizontal bars show the median value. The number of cells in each group are given on top. **(F-G)** Comparison of XP_50_ values estimated for the B6 and Cast chromosomes. Genes with significantly different silencing dynamics (ANOVA F test: BH correction, FDR<0.05) are colored in red and shown in (G). **(H)** K-means (K=4) clustering of genes according to their XP_50_ values on the B6 (left, 74 X-genes) and Cast chromosomes (right, 35 X-genes). Connecting bars in the center compare classification for genes that could be analyzed on both chromosomes (35 X-genes). Grey bars indicate genes that were assigned to the same cluster for the two alleles, light red indicates genes that were classified in neighboring clusters and dark red are genes that are part of more distant clusters.

To be able to compare silencing efficiency between alleles of individual genes, we developed an approach to quantify silencing of each gene relative to the rest of the chromosome. In this way we aimed at normalizing for the observed global differences in X-inactivation dynamics. To this end we quantified the “XCI progress” of each cell with monoallelic Xist expression, defined as the percentage of silencing of the inactive X chromosome (Fig. 5B, see computational methods for details). Although XCI progress correlated with the time point when cells were collected (Spearman’s correlation coefficient ρ =0.73, p<2.2*10^−16^) and with the pseudotime estimated based on the global transcriptome (Spearman’s correlation coefficient ρ =0.76, p<2.2*10^−16^) (Fig. 5C), XCI and differentiation appeared to be partially independent processes, as suggested previously (Chen et al., 2016). To quantify the silencing state for each gene relative to the rest of the chromosome, we grouped cells into 10 bins according to their XCI progress and calculated a lumped Xi:Xa ratio across cells in each bin for each gene (Fig. 5D, solid dots). We then fitted a log-linear model to the mean Xi:Xa ratio across bins, separately for cells silencing the B6 or Cast chromosome (Fig. 5D, lines). In this way we estimated an allele- and gene-specific XCI half time, corresponding to the global XCI progress where a gene was silenced by 50% (XP_50_) (Fig. 5D). To validate this approach we compared the XP_50_ values with results from previous studies, where genes had been classified according to their silencing dynamics on the B6 chromosome (or the closely related 129 allele), based on bulk sequencing of mature or nascent RNA (Barros de Andrade E Sousa et al., 2019; Marks et al., 2015). Moreover, we compared the estimated XP_50_ values with a previous classification based on scRNA-seq measurements in pre-implantation mouse embryos, where an imprinted form of XCI occurs, that had been performed for both alleles (Borensztein et al., 2017a). Our estimated XP_50_ values were in good agreement with these classifications (Fig. 5E).

In the next step we compared the estimated XP_50_ values between the two alleles for each gene (Fig. 5F). As expected the majority of genes exhibited similar silencing dynamics, since we had normalized for global silencing differences through the XCI progress approach. We found a subset of genes (Klhl13, Pir, Hprt), which were silenced with significantly different dynamics on the two alleles (ANOVA F test: BH correction, FDR<0.05, Fig. 5G). While Klhl13 appeared to escape on the Cast chromosome and was silenced on B6, Pir and Hprt were silenced more slowly on the B6 chromosome. Using k-means clustering we grouped all genes into 4 categories according to their silencing dynamics (fast, intermediate, slow, escape), separately for each allele (Fig. 5H). Here 3 out of 35 genes were assigned to the fast category on one allele and to the slow or escaping group on the other (Klhl13, Pir, Jade3), suggesting again the existence of allele-specific escape genes, such as Klhl13 and Pir. Taken together, our results show that overall, the Cast chromosome is silenced faster than the B6 allele. In addition, susceptibility to Xist-mediated gene silencing appears to be altered by genetic variation for a subset of genes. To further validate these findings we aimed to estimate allele-specific silencing dynamics with an orthogonal experimental approach.

### Experimental validation of allele-specific silencing dynamics

To assess XCI dynamics on the B6 and Cast chromosomes independently, we generated two mESC lines, where the X-inactivation center (Xic), which encompasses the *Xist* gene, was deleted on either one or the other allele, and named them TXΔXic_B6_ and TXΔXic_Cast_, respectively (Fig. 6A). To this end, a ∼800kb region around the *Xist* locus was deleted through CRISPR/Cas9-mediated genome editing (Fig. S5). Upon differentiation these cell lines underwent non-random XCI (Fig. 6A, Fig. S6A), allowing us to use bulk assays to measure gene silencing. We verified that both cell lines upregulated Xist with comparable dynamics and that the fraction of Xist-expressing cells was similar (Fig. S6B+C). To assess global XCI progression, we measured silencing dynamics for 5 genes (Renbp, Atrx, Cul4b, Prdx4, Rlim), for which our single-cell analysis had estimated comparable XP_50_ values for both alleles (Fig. 5F). We quantified the relative allelic expression over a 4-day differentiation time course through pyro-sequencing, which performs quantitative sequencing over individual SNPs on cDNA (Fig. S6D). Overall, the Xi:Xa ratio decreased more strongly in the TXΔXic_B6_ line compared to TXΔXic_Cast_ cells (Fig. 6B), confirming faster silencing on the Cast chromosome. In addition we tested all three genes that were found to be differentially silenced between the alleles in our scRNA-seq analysis (Fig. 5G). For Klhl13, which we had found to escape on the Cast chromosome, while being silenced on the B6 allele, we indeed observed a strong decrease of the Xi:Xa ratio on the B6 chromosome and even an increase in the line that silenced the Cast allele (Fig. 6C, Fig. S6E). The two other genes we tested, Pir and Hprt, had been found to be silenced faster on the Cast allele and we could indeed confirm that, at later time points, the Xi:Xa ratio appeared to be reduced more strongly in the line that inactivated the Cast chromosome (Fig. 6D, Fig. S6F). It must be noted, however, that we cannot distinguish at this point, whether these differences arise only from the overall increased efficiency of XCI on the Cast allele, or whether indeed additional gene-specific effects also contribute. Taken together, we could confirm faster silencing of the Cast allele in an independent experimental approach and we could clearly validate Klhl13 as an allele-specific escape gene.

**Figure 6.**
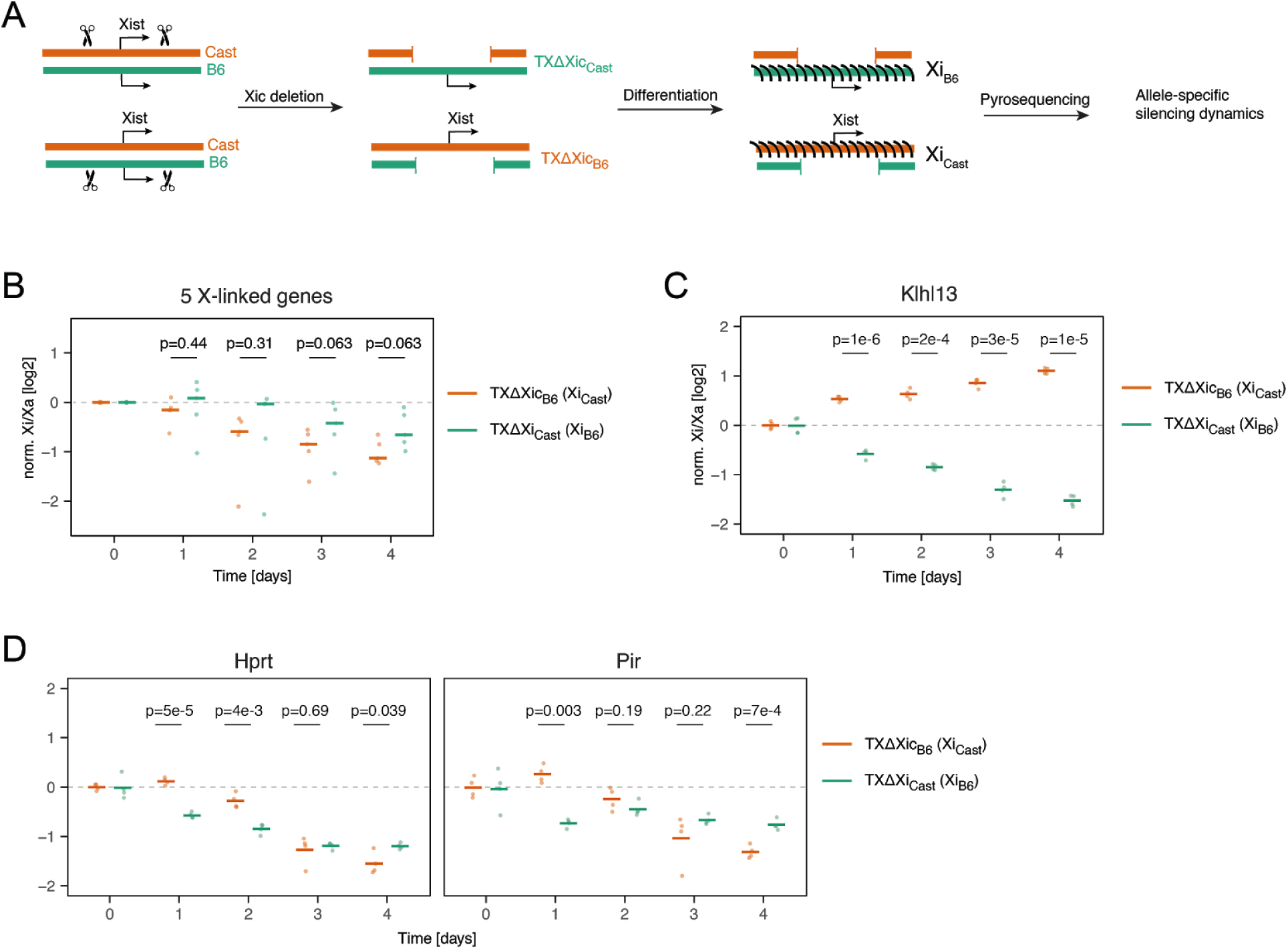
Experimental validation of differential silencing dynamics. **(A)** To measure allele-specific silencing dynamics, the X inactivation center (Xic), which contains the *Xist* locus, was deleted on either the Cast (top) or on the B6 allele (bottom). Upon differentiation, the entire cell population will thus initiate XCI on the B6 and Cast allele, respectively, allowing quantification of allele-specific silencing dynamics by pyrosequencing. **(B-D)** Xi:Xa expression ratio normalized to the pre-XCI state (day 0, dashed line). **(B)** Global silencing dynamics assessed for 5 genes that exhibited similar XP_50_ values on both alleles in the analysis in 5F (Renbp, Rlim, Prdx4, Cul4b, Atrx). The means (dots) of 4 independent biological replicates is shown, the horizontal bar indicates the median of all 5 genes. For each time point, the p-values were calculated using a Wilcoxon signed-rank two-sided test. **(C-D)** Allele-specific silencing dynamics for Klhl13 (C), which was predicted to escape at the Cast allele in 5F and for Pir and Hprt (D), which were predicted to be silenced more slowly on the B6 chromosome. Bars represent the mean of 4 biological replicates, dots the individual measurements. Additionally, p-values of a two-sample unpaired Student’s T two-sided test are shown.

## Discussion

We have profiled multiple steps governing the onset of random X inactivation with high temporal and allelic resolution through single-cell RNA sequencing. By adapting single-cell transcriptome analysis approaches, including recently developed concepts like RNA velocity, to an allele-specific process such as random XCI, we could draw a detailed picture of the different steps occurring at the onset of X inactivation. In this way we were able to answer several open questions by 1) dissecting the dynamics of XCI along developmental progression with allelic resolution and 2) quantifying the expression heterogeneity of the main genes involved in the XCI process, their regulatory relationships and their dynamics. Due to an efficient library preparation protocol and sufficient sequencing depth we could detect a median of ∼120.000 mRNA molecules per cell, which is significantly more than other UMI-based methods (Ziegenhain et al., 2017). This allowed us to quantify allelic expression for individual genes, including Xist and genes that are subject to XCI. We identified different Xist patterns at different stages of the XCI process and their associated gene silencing state. In addition, we exploited the heterogeneous nature of Xist upregulation to identify putative regulators of Xist. Finally, our allele-specific analysis revealed marked differences between the B6 and Cast X chromosomes, suggesting that cis-acting genetic variants modulate both, Xist expression and gene silencing dynamics.

We used a strand-specific, 3’-end counting scRNA-seq protocol. This enabled a detailed quantitative analysis of Xist expression, because it allows us to clearly distinguish Xist from its antisense transcript Tsix (Lee et al., 1999), in contrast to previous studies, where Xist had been analyzed in single cells, which had used unstranded full-length scRNA-seq techniques (Cheng et al., 2019; Chen et al., 2016). Although the protocol generally detected the 3’-end of most transcripts, since reverse transcription (RT) was initiated from the polyA tail, this was not the case for Xist, suggesting that Xist’s 3’-end is inaccessible during the RT reaction. This effect is not specific to the protocol used here, since we observed a similar pattern in data generated with CEL-seq2 (Hashimshony et al., 2016). Moreover, we and others have failed previously to amplify Xist in single-cell mRNA libraries, both in ES cells in vitro and in blastocyst stage embryos in vivo, which was at the time circumvented by addition of an Xist-specific RT primer (Borensztein et al., 2017b; Schulz et al., 2014). Xist’s 3’-end thus seems to be covered by a structure that can resist the mild denaturation conditions in scRNA-seq protocols. This poses a general question of how many other RNAs, maybe specifically nuclear lncRNAs might escape detection in scRNA-seq experiments.

Although reverse transcription could not be initiated from Xist’s polyA tail, a genomically encoded polyA stretch served as a priming site instead, allowing us to nevertheless quantify Xist expression in an allele-specific fashion. Such polyA stretches have also been proposed to underlie detection of intronic reads used for RNA velocity analyses in 3’-end scRNA-seq data sets (La Manno et al., 2018). At the early time points we identified a subset of cells that expressed Xist from both chromosomes, which constituted up to ∼25% of the population. We and others have previously observed such transient biallelic Xist upregulation by RNA FISH, both in differentiating mESCs and in mouse embryos in vivo (Guyochin et al., 2014; Mutzel et al., 2019; Sousa et al., 2018). We found that biallelic Xist upregulation actually initiated gene silencing and that it was resolved, when chromosome-wide silencing became visible, accompanied by a decrease in the number of Xist-expressing cells. These observations are in line with the idea we have previously proposed that complete silencing of an essential X-linked Xist activator might lead to Xist downregulation in cells where XCI has been initiated on both chromosomes (Mutzel et al., 2019). Accordingly, we find that Rnf12/Rlim, which functions as such an activator (Barakat et al., 2014; Jonkers et al., 2009), is indeed rapidly silenced, leading even to a strong negative correlation with Xist at day 2 (Fig. S3B). Our data thus lends strong support to the stochastic model of XCI, where initial Xist upregulation occurs independently on each allele with a low probability, and a silencing-mediated negative feedback loop then ensures that only a monoallelic state can be maintained (Monkhorst et al., 2008).

The stochastic nature of Xist upregulation was also supported by our finding that Xist expression was heterogeneous across cells and the correlation with its regulators was rather weak (absolute correlation coefficient <0.32). While two previous studies had attempted to identify XCI regulators by analyzing correlation with X-chromosomal gene activity (Chen et al., 2016; Mohammed et al., 2017), our analysis allows identification of the initial triggers of Xist upregulation even before gene silencing is initiated. When Xist was first upregulated, at day 1 of differentiation, very few differences could be detected between the transcriptomes of cells expressing Xist and those that had not yet initiated XCI. A notable exception was Nanog, which was consistently expressed at higher levels in cells with no or low Xist expression. This observation thus suggests that Xist upregulation is directly triggered by Nanog downregulation, independently of Oct4 and Sox2, which have also been proposed as developmental Xist regulators (Donohoe et al., 2009; Navarro et al., 2008). In addition, we found another putative Xist regulator, Snhg1, which has not yet been implicated in XCI and which might function as an activator of Xist. This poorly-studied lncRNA, which has been suggested to function through binding to Igf2bp1 (Huang et al., 2020), which in turn modulates protein stability of a series of target genes, including pluripotency factors (Bi et al., 2020), warrants further investigation in the future.

The present study was performed in a highly polymorphic cell line to enable an allele-specific analysis, which is required to quantify gene expression from each X chromosome individually. Interestingly we found significant differences between the two X chromosomes, suggesting that cis-acting genetic variation modulates the X inactivation process at multiple levels. First, we observed the long-known Xce effect leading to preferential inactivation of the B6 X chromosome, which has been mapped to a large genomic region centromeric to *Xist* (Cattanach et al., 1969; Galupa and Heard, 2015; Plenge et al., 2000). In addition we observed that the B6 X chromosome also produced more Xist transcripts than the Cast allele. Surprisingly, this did not enhance silencing, rather the opposite with more efficient silencing being observed for the Cast allele. This observation suggests that polymorphisms within the Xist RNA might silencing efficiency of Xist in the B6 strain, for example by altering binding affinity for silencing factors such as Spen (Brockdorff et al., 2020). An intriguing hypothesis is that such reduced silencing efficiency might have been evolutionarily compensated by faster and stronger Xist upregulation at the B6 X chromosome, ultimately leading to the long-known Xce effect in hybrid mouse embryos.

Since XCI is a chromosome-wide process it can be quantified by scRNA-seq in a fairly robust manner despite the limited sensitivity of the method. Due to the high quality of the dataset we have generated we could in addition compare silencing of individual genes between alleles. We found that a subset of genes were silenced with different dynamics, in agreement with a similar observation made during imprinted XCI in preimplantation embryos (Borensztein et al., 2017a). Although substantial heterogeneity with respect to escape from XCI has been reported in single human fibroblasts (Garieri et al., 2018), no systematic analysis of putative genetic determinants of this heterogeneity has been performed to date. An extreme scenario with complete escape from XCI specifically on the Cast chromosome was observed for Klhl13 and confirmed in an independent experiment. Interestingly, we have recently identified Klhl13 as an X-linked differentiation inhibitor (Genolet et al., 2020). Its silencing might help to release a differentiation block imposed by the presence of two active X chromosomes (Schulz et al., 2014; Song et al., 2019). Escape of Klhl13 in the Cast strain might therefore have evolved to compensate for the overall faster silencing of the X chromosome in that genetic background, which might otherwise result in a too fast release of the differentiation block. In the future, strain-specific escapees such as Klhl13 will potentially allow us to identify escape-promoting cis-acting genetic elements to better understand the principles that can protect a gene from inactivation. Escape mechanisms are a key unanswered question in XCI research and will, once elucidated, be an important contribution to understanding epigenetic gene regulation.

The strategies we have developed here to profile X inactivation in the endogenous setting of random XCI will be valuable to investigate XCI status in other contexts, in particular those that are not amenable to genetic engineering. For example, we will now be able to assess XCI and escape in primary human cells, making use of naturally occurring genetic variation. This will allow us to investigate the stability of silencing on the inactive X in different tissues and cell types as well as tissue-specific escape for individual genes. Such information will be indispensable to understand how escape genes contribute to sexual dimorphic traits and to differential disease susceptiblity between the sexes such as the higher prevalence of autoimmune diseases in women (Libert et al., 2010).

## Methods

### Experimental Procedures

#### Cell lines

The female TX1072 cell line (clone A3) is a F1 hybrid ESC line derived from a cross between the 57BL/6 (B6) and CAST/EiJ (Cast) mouse strains that carries a doxycycline-responsive promoter in front of the *Xist* gene on the B6 chromosome and an rtTA insertion in the *Rosa26* locus (Schulz et al., 2014). TxdXic_A1 and TxdXic_A6 lines were generated by deleting a 773 kb region around the *Xist* locus. TxdXic_A1 carries the deletion on the B6 chromosome (chrX:103,182,701-103,955,531, mm10) and TxdXic_A6 on the Cast allele (chrX:103,182,257-103,955,698, mm10). To generate these cell lines, a total of 4 guides flanking the region to be deleted (2 on each side) were cloned into the pX459-v2 vector (Addgene #62988) (Ran et al., 2013). Four separate transfections were performed, each combining one guide upstream and downstream of the targeted region and a single-stranded repair oligo with 50bp homology flanking each cut site (IDT). Further details and sequences are provided in Supplemental Table S4. In each transfection, 2*10^6^ TX1072 mESCs were electroporated with 2µg of each guide plasmid and 30pmol of the single stranded repair oligo using the P3 Primary Cell 4D-Nucleofector X Kit (V4XP-3024) with the Amaxa 4D Nucleofector system (Lonza, program CP-106) and plated on gelatin-coated 10cm dishes. Starting 30h after transfection, cells were selected with puromycin (1ng/µl) for 16h. After 4 days equal cell numbers from all 4 transfections were pooled and plated at a density of 2-3 cells/well in gelatin-coated 96-well plates. Genomic DNA was extracted and screened for the presence of the deletion with primers VM21 and VM24. Cells from wells showing the presence of the deletion were seeded at clonal density (100 cells/cm^2^) in 10cm plates. Individual clones were picked and screened for the presence of the deletion and the wild type allele and the absence of inversions or duplications of the targeted region. The genotyping strategy is shown in Supplemental Figure S5. For all PCR reactions the Phusion HotStart Flex DNA Polymerase (NEB) was used with an annealing temperature of 55°C, an elongation time of 45sec and 35 cycles. Primer sequences are listed in Supp. Table S4. To determine which allele carried the deletion, the PCR products were sequenced and annotated SNPs between Cast and B6 alleles were used to assign the deletion and the wild type allele. Correct karyotype of clones was verified with metaphase spreads.

#### mESC culture and differentiation

Cells were grown on gelatin-coated flasks in serum-containing ES cell medium (DMEM (Sigma), 15% ESC-grade FBS (Gibco), 0.1mM β-mercaptoethanol, 1000 U/ml leukemia inhibitory factor (LIF, Millipore)), supplemented with 2i (3 μM Gsk3 inhibitor CT-99021, 1 μM MEK inhibitor PD0325901, Axon). Differentiation was induced by 2i/LIF withdrawal in DMEM supplemented with 10% FBS and 0.1mM β-mercaptoethanol at a density of 1.5*10^4^cells/cm^2^ in fibronectin-coated (10 μg/ml) tissue culture plates.

#### scRNA-seq

Single-cell RNA-seq libraries were prepared with the C1-HT mRNA-seq v2 protocol according to the manufacturer’s recommendations (Fluidigm). Cells were rinsed thoroughly with PBS, trypsinized for 7 minutes and resuspended in the respective growth medium at a concentration of 400 cells/μl. 30 μl cell suspension was diluted with 20 μl of suspension reagent (Fluidigm) and 10 μl of the dilution was loaded into one compartment of a Single-Cell mRNA Seq HT integrated fluidic circuit (IFC) 10-17 µm. A different cell type was loaded into the other compartment, which is not analyzed in this study. Cell viability staining was performed on the IFC using the LIVE/DEAD viability/Cytotoxicity Kit (Thermofisher) with 1 μM Ethidium and 0.05 μM Calcein. IFC loading and life/dead staining was analyzed with automated image acquisition using a Zeiss CellDiscoverer microscope (Zeiss) with a 20x objective. During the lysis step ERCC Spike-in Mix 1 (Thermofisher) was added with a final dilution of 1:200.000. Lysis, reverse transcription and cDNA amplification was performed on the C1 machine. cDNA pools were quantified by Qubit and Bioanalyzer HS. Around 2.25 ng of each pool were subjected to tagmentation and library preparation using the NexteraXT library preparation kit according to the C1-HT protocol. All pools were mixed in equal proportions and quantified with KAPA Library Quant-Kit. The libraries were sequenced on a HiSeq2500 instrument (Illumina) with asymmetric read length, either in High Output (Read1: 13bp, Index read: 8pb, Read2: 48bp) or in Rapid Run mode (Read1: 16bp, Index read: 8pb, Read2: 36bp), with 10pM loading concentration and 5% PhiX.

#### RNA extraction, reverse transcription, qPCR

For gene expression profiling, cells were lysed directly in the plate by adding 1 ml of Trizol (Invitrogen). RNA was isolated using the Direct-Zol RNA Miniprep Kit (Zymo Research) following the manufacturer’s instructions, and DNase digest was performed using Turbo DNA free kit (Ambion). 1 µg RNA was reverse transcribed using Superscript III Reverse Transcriptase (Invitrogen) and expression levels were quantified using Power SYBR™ Green PCR Master Mix (4368702, Thermo Fisher) normalizing to Rrm2 and Arpo. Primers used are listed in Supplemental Table S4.

#### Pyrosequencing

To quantify relative allelic expression for individual genes, an amplicon containing a SNP was amplified by PCR from cDNA using GoTaq Flexi G2 (Promega) with 2.5 mM MgCl2 or Hot Star Taq (Qiagen) for 40 cycles. The PCR product was sequenced using the Pyromark Q24 system (Qiagen). Assay details are given in Supplemental Table S4.

#### RNA FISH

RNA FISH on Xist and another X-linked gene, Huwe1 was performed using Stellaris FISH probes (Biosearch Technologies). Probe details can be found in Supplemental Table S4. Cells were dissociated using Accutase (Invitrogen) and adsorbed onto coverslips (#1.5, 1 mm) coated with Poly-L-Lysine (Sigma) for 5 min. Cells were fixed with 3% paraformaldehyde in PBS for 10 min at room temperature (18–24°C) and permeabilized for 5 min on ice in PBS containing 0.5%Triton X-100 and 2 mM Ribonucleoside Vanadyl complex (New England Biolabs). Coverslips were preserved in 70% EtOH at -20°C. Prior to FISH, coverslips were incubated for 5 minutes in Stellaris RNA FISH Wash Buffer A (Biosearch Technologies), followed by hybridization overnight at 37°C with 250 nM of each FISH probe in 50 μl Stellaris RNA FISH Hybridization Buffer (Biosearch Technologies) containing 10% formamide. Coverslips were washed twice for 30 min at 37°C with Stellaris RNA FISH Wash Buffer A (Biosearch Technologies), with 0.2 mg/ml Dapi being added to the second wash. Prior to mounting with Vectashield mounting medium coverslips were washed with 2xSSC at room temperature for 5 minutes. Images were acquired using a widefield Z1 Observer microscope (Zeiss) using a 100x objective. The intronic signal of Huwe1 was used in combination with Xist to estimate the percentage of XO cells in the population, which was maximally 5%.

### Computational Methods

#### Alignment and gene quantification

The C1-HT protocol uses a dual barcoding strategy, where read R1 contains a custom barcode (position 1-6, cell barcode) and a unique molecular identifier (position 7-11), read R2 maps to the cDNA sequence and read R3 contains a Nextera (row) barcode. After demultiplexing using the Nextera barcode, read R2 was aligned with STAR (v.2.5.2b) (Dobin et al., 2013) to the mouse genome genome (mm10) and all 96 ERCC spike-in sequences, allowing for unique alignments with a maximum of two mismatches. SNPsplit (v.0.3.2) (Krueger and Andrews, 2016) was used to N-mask the genome using high confidence SNPs, confirmed to be present in the TX1072 cell line based on ChIP input data (Barros de Andrade E Sousa et al., 2019).

The Drop-seq pipeline (Macosko et al., 2015) was used to extract both cell barcode and UMI from read R1 and to quantify gene expression. In detail, reads were demultiplexed using the cell barcode, and molecules per gene were counted as the number of unique exon-overlapping UMI barcodes using the mm10 Ensembl gene annotation. For RNA velocity analyses unique UMI barcodes aligning only to exonic regions (spliced) or overlapping with intronic regions (unspliced) were used. For allele-specific (AS) quantification each read overlapping a SNP position was assigned to its parental genome using SNPsplit (v.0.3.2) (Krueger and Andrews, 2016) and the number of unique UMI barcodes assigned to either parental genome were counted for each gene (Supplemental Figure S1A,D).

In the following, we denote the number of exon-overlapping, spliced and unspliced not-AS molecules for each gene *g* and each cell *c*, as *X*_*g,c*_, *S*_*g,c*_ and *U*_*g,c*_, respectively. Similarly, the estimated number of exon-overlapping, spliced and unspliced AS molecules for both alleles, *Cast* and *B6*, for each gene *g* and cell *c*, are denoted with 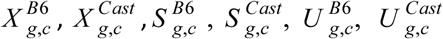.

#### Cell filtering

To remove empty wells, dead and low-quality cells, several cell filtering steps were performed (Supplemental Table S1). Capture sites without a cell or with multiple cells were identified by manual inspection based on brightfield and fluorescence imaging (live/dead stain) of the IFC and removed from subsequent analyses. Moreover, dead cells, cells with low number of reads, low number of transcripts or low number of expressed genes, as well as cells with high percentage of mitochondrial DNA or ERCC spike-in reads were removed from the analysis (Supplemental Figure S1B). To identify dead cells the fluorescent signal of the dead stain (Ethidium) was quantified as the integrated intensity within a rectangle of constant size, manually centered around each capture site using ZEN v2.3 software (Zeiss). Thresholds were set as the 3-fold median-absolute-deviation (MAD) above or below the median of the respective variable x (e.g. number of reads) for the removal of cells with high and low signal, respectively: *median*(*x*) ± 3 · *median*(| *x* −*median*(*x*) |). Finally, cells that did not express Xist and where >80% of X-linked reads mapped to the same allele were assumed to have lost one X chromosome (XO). To this end, the X-chromosomal ratio was defined for each cell c as:

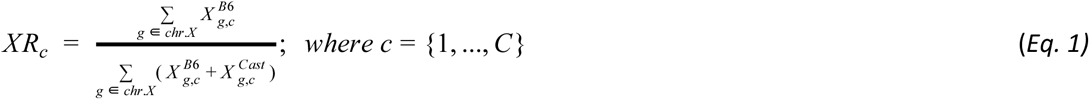

Cells with *XR*_*c*_ ≥ 0.8 or *XR*_*c*_ ≤ 0.2 were assumed to be XO and were removed from the analysis (Supplemental Figure S1C).

#### Gene filtering

In order to analyze only genes with sufficient detection rate across cells, genes that were detected in <20% of all cells were excluded from the analysis. Gene filtering was performed separately for the AS and not-AS analysis. Furthermore, genes with a strong skewing towards a single allele where >90% of reads detected in all cells mapped to the same allele were excluded from the AS and not-AS analysis (Supplemental Table S1).

#### Normalization of read counts

In order to correct for cell-specific biases, prior to downstream analyses, gene count normalization was performed based on autosomal genes only, using the *scran* (v.1.12.1) R package (Lun et al., 2016). Cell clusters were defined independently for each time point through the *computeSumFactors* function and the *clusters* parameter. Cluster-based scaling factors were then deconvoluted into cell-specific factors (Supplemental Table S2). The cell-specific scaling factors (*sf* _*c*_; *c* = 1, …, *C*) derived from the not-AS count matrix (*X*) were used to compute the normalized counts per million values for the *g*-th gene and *c*-th cell (*CPM* _*g, c*_) of the AS and not-AS count matrices:

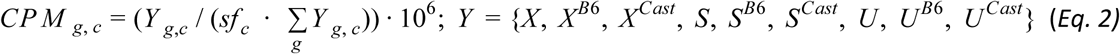

#### Differential expression analyses

Cells were clustered with K-means clustering (K=7, *kmeans* function from the *stats* R package) based on the logarithm of Xist CPM expression *log*_10_(*CPM*_*Xist, c*_ + 1). Cells belonging to the top three K-means groups, i.e. high expression of Xist, were classified as Xist-high, while cells belonging to the bottom 3 K-means groups, i.e. low expression of Xist were classified as Xist-low (Supplemental Table S2). K was set in a way to minimize the within-cluster sum of squares value, while ensuring a minimum number of 50 cells in the Xist-high and Xist-low groups at each time point of differentiation. The set of differentially expressed genes (DEG) between Xist-high and Xist-low cells were identified at each time point through the MAST differential expression analysis method, using the *zlm* function from the *MAST* (v.1.10.0) R package (Finak et al., 2015). In brief, a two-part generalized Hurdle model was fitted to the gene-wise normalized not-AS CPM expression values. We set dummy variables to represent the two cell groups and the proportion of detected genes (UMI count > 0) per cell as model predictors. The significance of each gene between the two groups of cells was then assessed by a χ^2^ likelihood ratio test. The genes with a Benjamini-Hochberg adjusted p-value smaller than the nominal FDR=0.05 were deemed as significantly differentially expressed between the two cell groups (Benjamini and Hochberg, 1995). Notably, the MAST method was only applied to compare groups containing at least 10 cells. For this reason, the cells at day 0 were excluded from the analysis. The results are provided in Supplemental Table S3.

#### X:A ratio

We computed the ratio between X-linked and autosomal spliced gene expression similarly to Borensztein et al. (Borensztein et al., 2017a). Briefly, in order to account for the larger number of autosomal genes compared to X-linked genes (8880 and 374 genes respectively) we defined B=1000 bootstrap samples of autosomal genes of the same size of the set of X-linked genes. For each cell and bootstrap sample we computed the ratio between the average X-linked and average autosomal gene expression, and estimated the X:A ratio for each cell as the median across all B computed bootstrap ratios.

#### Pseudotime analysis

Pseudotime trajectories allow to order single cells according to a biological process of interest and identify key paths or branches corresponding to alternative cellular states of process outcomes. We analyzed single cell trajectories using the Monocole2 algorithm (*monocole* R package v.2.12.0) (Qiu et al., 2017b, 2017a; Trapnell et al., 2014). Starting from high dimensional data, it projects cells into a lower dimensional space by constructing a principal graph and ordering them according to a pseudotime trajectory. Similarly to Cacchiarelli et al. (Cacchiarelli et al., 2018), a set of ordering genes was defined as the 500 most differentially expressed genes over time, identified using the *differentialGeneTest R* function. The DDRTree method was used to project the cells into a two-dimensional space based on the expression of the selected genes, and simultaneously learn a graph structure into this space (Qiu et al., 2017b). Pseudotime values were then estimated as the distance of each cell from the root of the graph, which is defined as the state with the highest number of day 0 cells. Finally, the cell-wise scaled pseudotime was computed dividing the estimated values by the maximum across all cells (Supplemental Table S2).

#### Dimensionality reduction (UMAP)

After the log-transformation of the CPM-normalized not-AS gene expression values, the most variable features were identified as those genes with the 500 highest variances across all cells and time points. The number of features was further reduced through a principal component analysis (PCA) based on the centered expression levels of the selected genes using the *pca* R package (v.1.76.0). The top-50 principal components (PCs) were provided as input to the UMAP dimensionality reduction method (McInnes et al., 2018) to further reduce dimensionality and visualize cells in a two-dimensional space, using the *umap* R package (v.0.2.3.1).

#### RNA velocity analysis

We performed RNA-velocity analyses based on the not-AS spliced and unspliced count matrices (*S* and *U*, respectively), removing the genes showing low average spliced 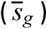 or unspliced 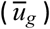 expression across all time points and cells (namely, 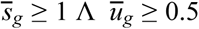) (La Manno et al., 2018). RNA velocities were then calculated (using the *gene*.*relative*.*velocity*.*estimates* function from the *velocyto*.*R* (v.0.6) R package) setting the cell neighbourhood size to *kCells = 20*, and the remaining parameters to their default values. The estimated velocities were then projected onto the UMAP embedding and locally summarized through a vector field representation of single cell velocities.

#### Allele-specific RNA velocity

To perform RNA-velocity analysis for X-linked genes in an allele-specific fashion, we first selected genes with sufficient counts by adding up the AS count matrices (*S* ^*AS*^ = *S* ^*B*6^ + *S* ^*Cast*^, and *U* ^*AS*^ = *U* ^*B*6^ + *U* ^*Cast*^) and removing genes with low average spliced or unspliced expression across all time points and cells (namely 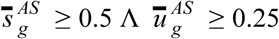). To define a neighbourhood of cells with similar XCI level, required to fit the velocity model, we first computed the B6-ratio matrix for each X-linked gene and cell as:

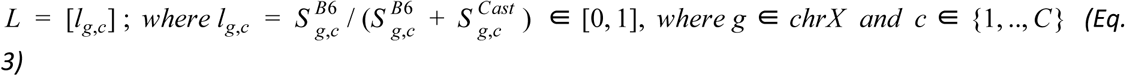

Based on the B6-ratio vectors for each cell, we computed pairwise cell-cell Pearson correlation coefficients and converted them to a distance matrix, where each element represents the distance between the i-th and j-th cells:

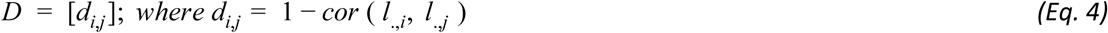

RNA velocities were computed with setting the neighbourhood of each cell to the 20 cells with smallest distance (kCells = 20), and the remaining parameters to their default values. The neighbourhood of each cell was restricted to the cells sequenced at the same time point. The cell embedding was defined by the first two principal components computed on the *L* matrix, and the estimated velocities were first projected onto this space and then summarized by a vector field.

#### Binned silencing efficiency analysis

In order to estimate relative gene silencing dynamics for each gene, we first computed the overall XCI progress (XP) for each cell *c* that expressed Xist monoallelically:

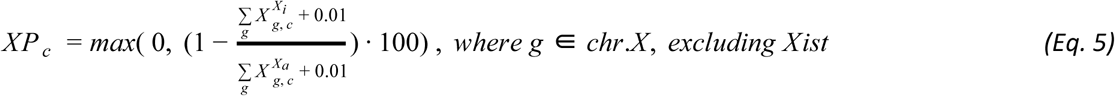

where Xi refers to the Xist-expressing chromosome and Xa to the Xist-negative allele (Supplemental Table S2). XP is a proxy for the extent of inactivation that has already occurred on the X chromosome, excluding Xist. Intuitively, a high *XP*_*c*_ value indicates that the cell has already silenced a substantial number of X-linked genes, while a value proximal to zero indicates that the two alleles have similar gene expression levels. The subsequent analysis was then restricted to cells that have initiated the XCI process, hence cells with *XP*_*c*_ ≥ 10%. In order to fit a model to the expression of each X-linked gene over XCI progress, cells were partitioned into 10 equally sized bins. For the *b*-th bin (b=1,..,10), the binned values *XP*_*b*_ (*XP*_*b*_ *with b* = 1, .., 10) were computed again, similarly to eq. 5, grouping together cells belonging to the same bin. The expression ratio *r*_*g,b*_ was then computed for each X-linked gene in each bin as:

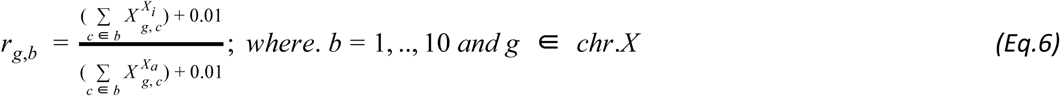

For each gene and allele, the analysis was restricted to the bins containing a minimum of 5 cells and a total of 25 AS counts 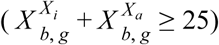 and analysis was restricted to genes with a minimum of 5 valid bins. To account for basal expression skewing due to genetic variation, *r*_*g,b*_ was normalized to *r*_*g,baseline*_ in Xist-negative cells at day 0 with 0.4 ≤ *XR*_*c*_ ≤ 0.6, resulting in

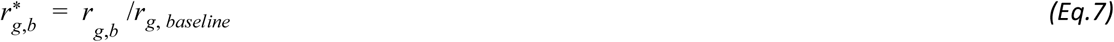

To test for differential silencing between alleles, for each gene two log-linear models were compared: the simpler model assumes a single trend for the two alleles (namely, β_1_), while the more complex one accounts for allele-specific slopes and models one trend per allele (namely, β_1_ and β_1_ + β_2_):

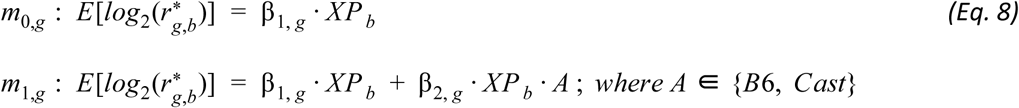

The significance of the β_2, *g*_ parameter was then assessed through an ANOVA F test (*H*_0_: β_2, *g*_ = 0). Any gene with an adjusted p-value (BH correction) smaller than 0.05 was deemed as differentially silenced between the two alleles. The gene-wise silencing half-times (*XP* _50, *g*_) were then computed as the XP value corresponding to a 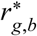 value of 0.5. The *XP* _50, *g*_ values greater than 100 were set equal to 100. To classify genes according to their allele-specific silencing dynamics (early, intermediate, late, escaping XCI) k-means clustering (K=4) was performed on the *XP*_50, *g*_ values of each allele separately (Supplemental Table S2).

#### Pyrosequencing data analysis

Let 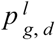 represent the percentage of B6-molecules observed for the *g* – *th* gene (Klhl13, Pir, Hprt, Rlim, Atrx, Renbp, Cul4b, Prdx4), and *l* – *th* deletion line (TXΔXic_B6_, TXΔXic_Cast_), after *d* days of differentiation. For each gene, day and cell line, we computed the ratio of molecules derived from the inactive allele as:

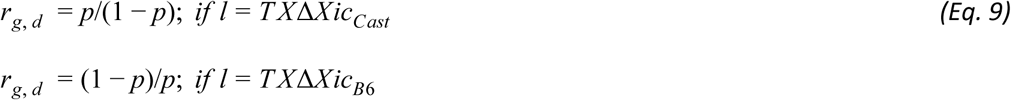

Aiming to account for baseline AS detection skewing, the above ratios were normalized to the day 0 gene-wise average ratios across replicates (namely, 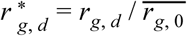). For each gene deemed as differentially silenced between the two alleles (Klhl13, Pir, Hprt), the normalized ratios observed in the two deletion lines were compared at each time point through an unpaired t-test statistic. The average normalized ratio of 5 not differentially silenced genes (Rlim, Renbp, Cul4b, Prdx4, Atrx) was computed for each time point and cell line, and the difference in silencing efficiency between the two deletion lines was tested at each time point through a Wilcoxon signed-rank test.

## Data and code availability

ScRNA-seq data generated during this study are available via GEO with identifier GSE151009 as raw fastq files and as the unfiltered not-AS and AS count tables together with the list of SNPs used for allele-specific analysis. The analysis code is deposited at https://github.com/EddaSchulz/Pacini_paper.

## Author Contribution

Edda G. Schulz, Annalisa Marsico and Guido Pacini conceived the present work. Ilona Dunkel and Norbert Mages performed the experiment with input from Edda Schulz and Bernd Timmermann. Guido Pacini performed all analyses and Verena Mutzel generated the ΔXic cell lines. Edda G. Schulz and Annalisa Marsico wrote the manuscript with input from Guido Pacini.

## Acknowledgements

We would like to thank Maud Borensztein and Marc Friedlander for critical feedback on the manuscript; Thorsten Mielke and Beatrix Fauler for imaging support; Sven Klages for data pre-processing and the IT service at the MPI for Molecular Genetics for maintenance and support of a computing cluster. This work was supported by the Max-Planck Research Group Leader program, E:bio Module III—Xnet grant (BMBF 031L0072) and Human Frontiers Science Program (CDA-00064/2018) to E.G.S. G.P. was part of the IMPRS for Biology and Computing.

---

**Supplemental Figure S1.**
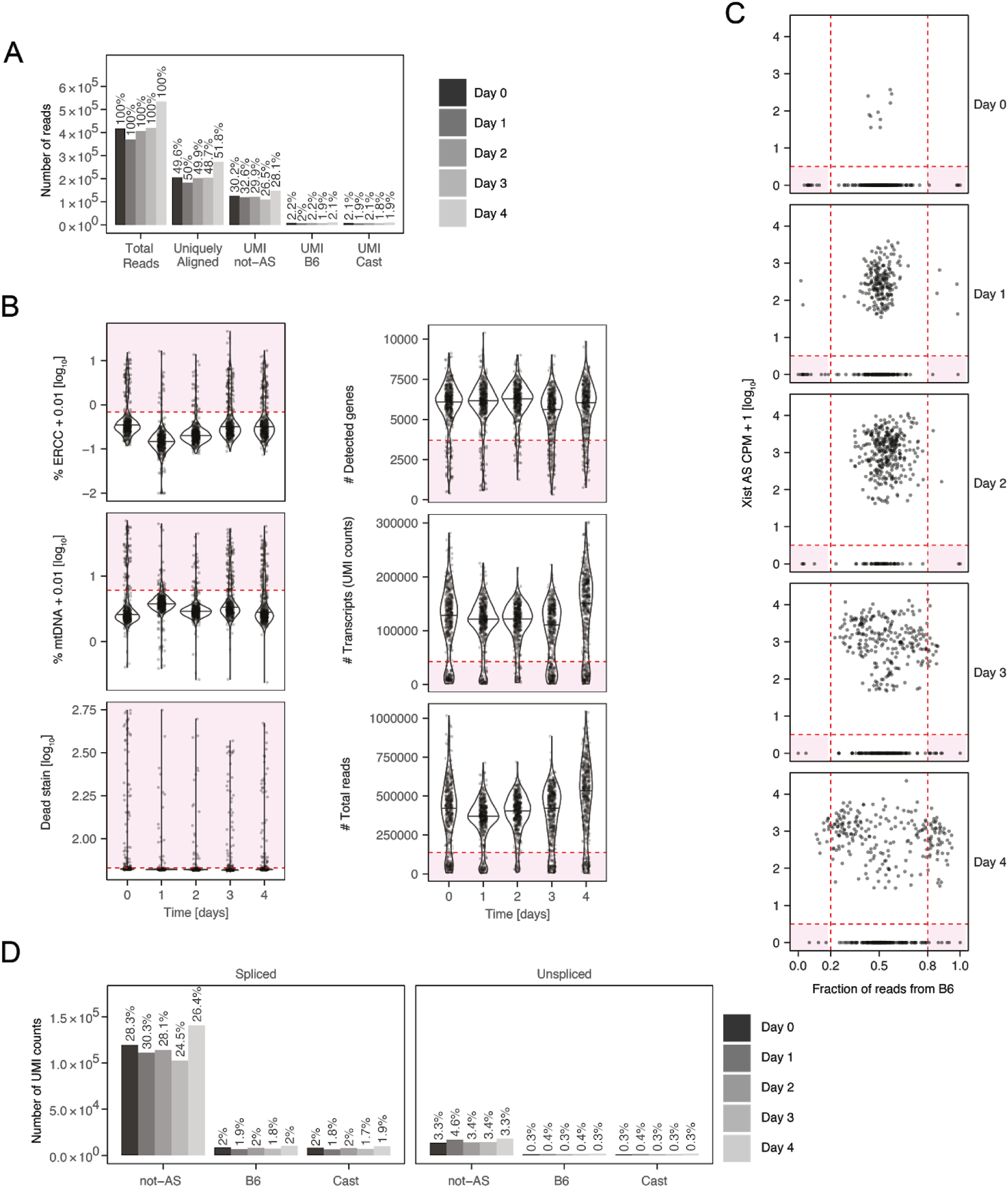
Data pre-processing. **(A)** Alignment statistics, showing the median number of reads across all sequenced cells. **(B)** Violin plots of the parameters used for cell filtering. Cells falling in the shaded areas were excluded from the analyses. **(C)** Putative XO cells were identified as AS Xist-negative cells, where >80% of X-chromosomal reads mapped to the same allele (shaded areas), and were excluded from the analyses. **(D)** Alignment statistics as in (A) for the spliced and unspliced quantification used for RNA velocity analysis.

**Supplemental Figure S2:**
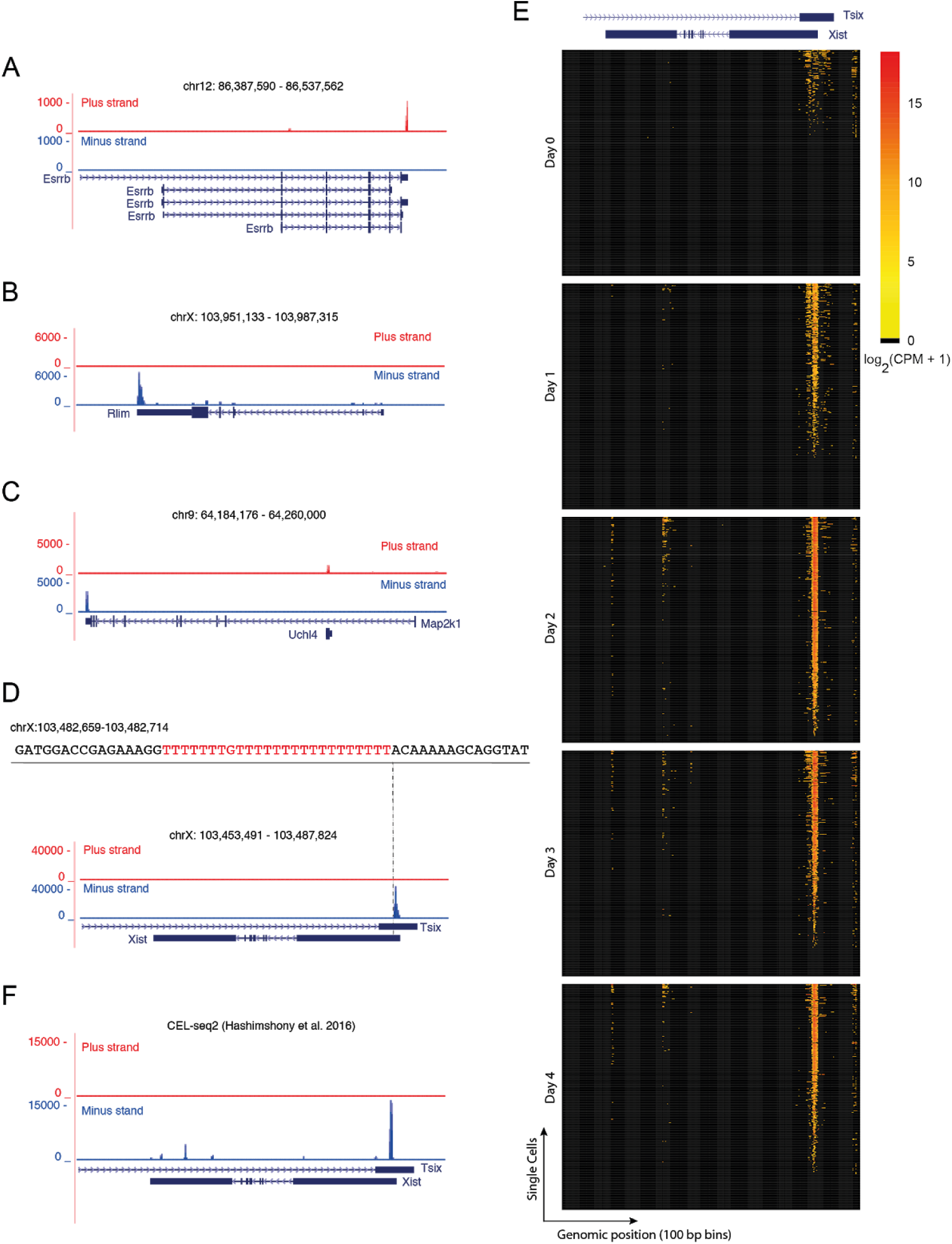
Unusual alignment pattern across the *Xist* gene. **(A-D)** Composite tracks across all cells show read distributions for three representative control genes (A-C) and Xist (D). In (D) a sequence segment next to the region to which most reads map is shown in detail. A polyT stretch that will generate a genomically encoded polyA stretch in the Xist RNA appeared to prime the reverse transcription reaction during library preparation. **(E)** Heatmaps describing the 100bp-binned read coverage across the *Xist* gene on the minus strand (chrX: 103,460,366 - 103,483,254) for all cells (rows), separately ordered for each time point by the total number of reads mapping to Xist. **(F)** Composite track around Xist as in (D), for a previously published scRNA-seq data set of murine fibroblasts using the CEL-seq2 protocol(Hashimshony et al., 2016).

**Supplemental Figure S3:**
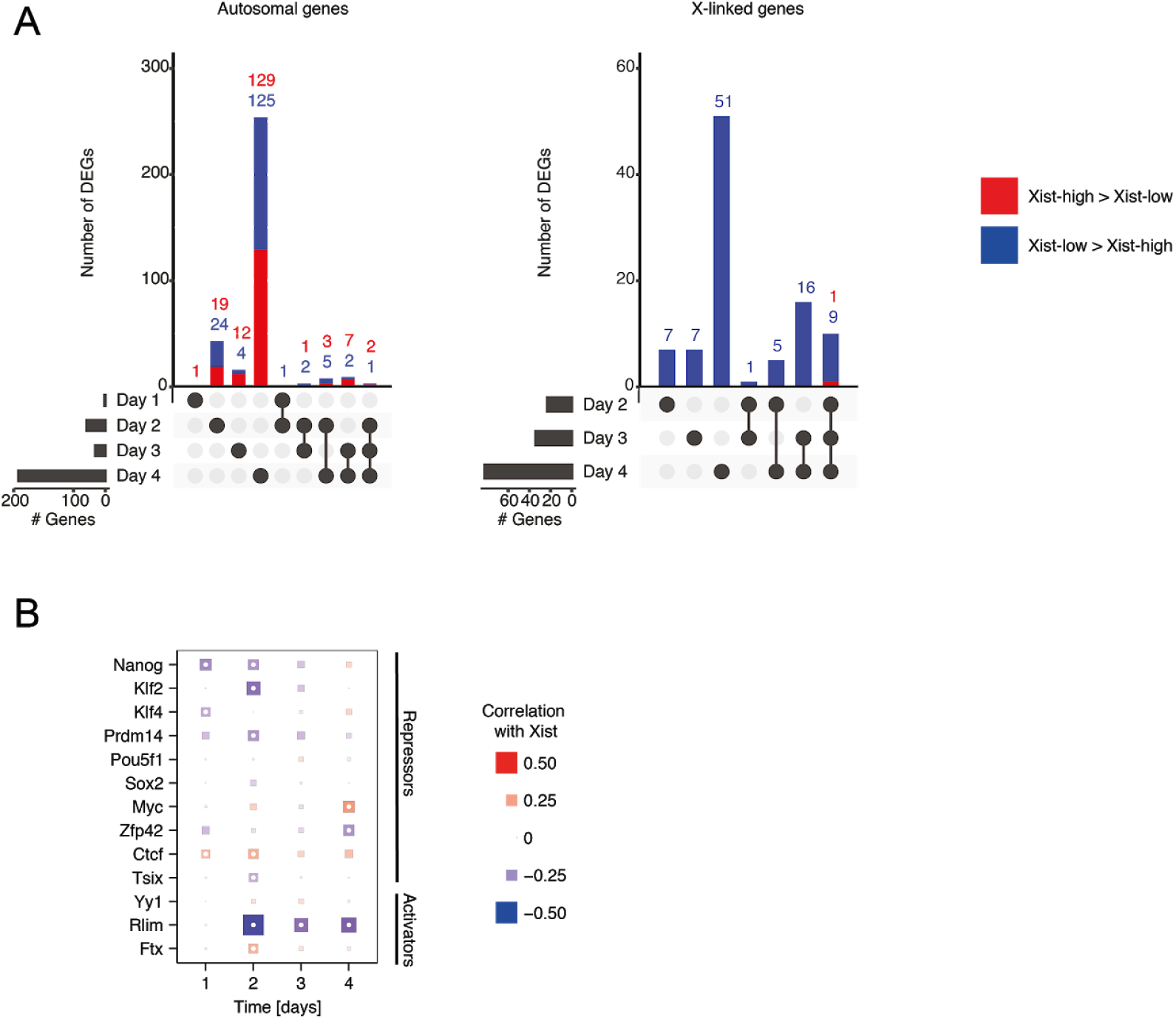
Identification of putative Xist regulators. **(A)** Overlap between differentially expressed genes (DEGs) as in Fig. 3B (FDR<0.05) between Xist-high and Xist-low cells, across different time points for autosomal (left) and X-linked (right) genes. Xist itself was excluded from the analysis. Horizontal bars indicate the total number of DEGs at each time point, and vertical bars describe the overlap between the DEGs detected at different time points as indicated below the plot. **(B)** Spearman’s correlation coefficients with Xist for a series of previously proposed Xist regulators, all of which, except Jpx, could be efficiently detected in the data set. Size and color indicate the correlation coefficient as indicated. White dots represent significant correlation (Spearman correlation test and BH correction, FDR<0.05). Negative correlation was found for the pluripotency factors Nanog and also Klf4 already at day 1 and with Nanog, Prdm14 and Klf2 at day 2, in agreement with their proposed role as Xist repressors(Gillich et al., 2012; Navarro et al., 2008). For two cis-acting non-coding transcripts, Xist’s repressive antisense transcript Tsix and the Xist activator Ftx, a significant, albeit low, negative and positive correlation, respectively, was detected. A very strong negative correlation was found for the well studied X-linked Xist activator Rlim (Rnf12), probably due to rapid silencing of this gene(Jonkers et al., 2009). Unexpectedly, a positive correlation was found for CTCF, which had previously been suggested to repress Xist(Sun et al., 2013).

**Supplemental Figure S4:**
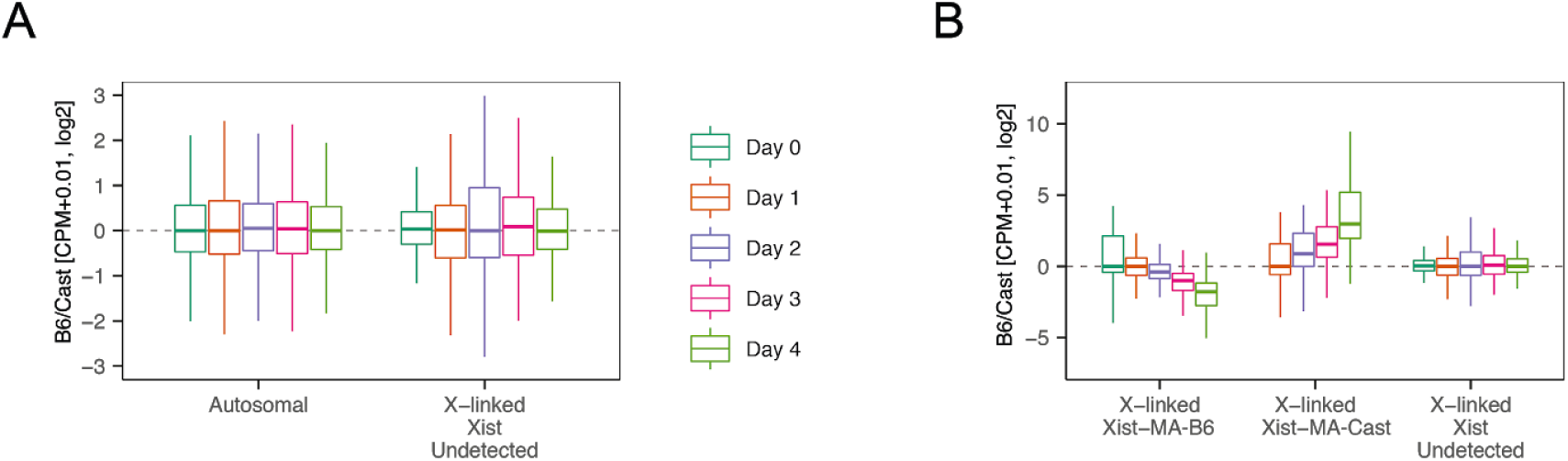
Equal detection rates for B6 and Cast chromosomes. **(A-B)** Box plots showing the distribution of the B6-to-Cast ratio of all autosomal and X-linked genes as indicated, calculated by pooling all cells (for autosomal genes) and the indicated cell subsets for X-linked genes.

**Supplemental Figure S5:**
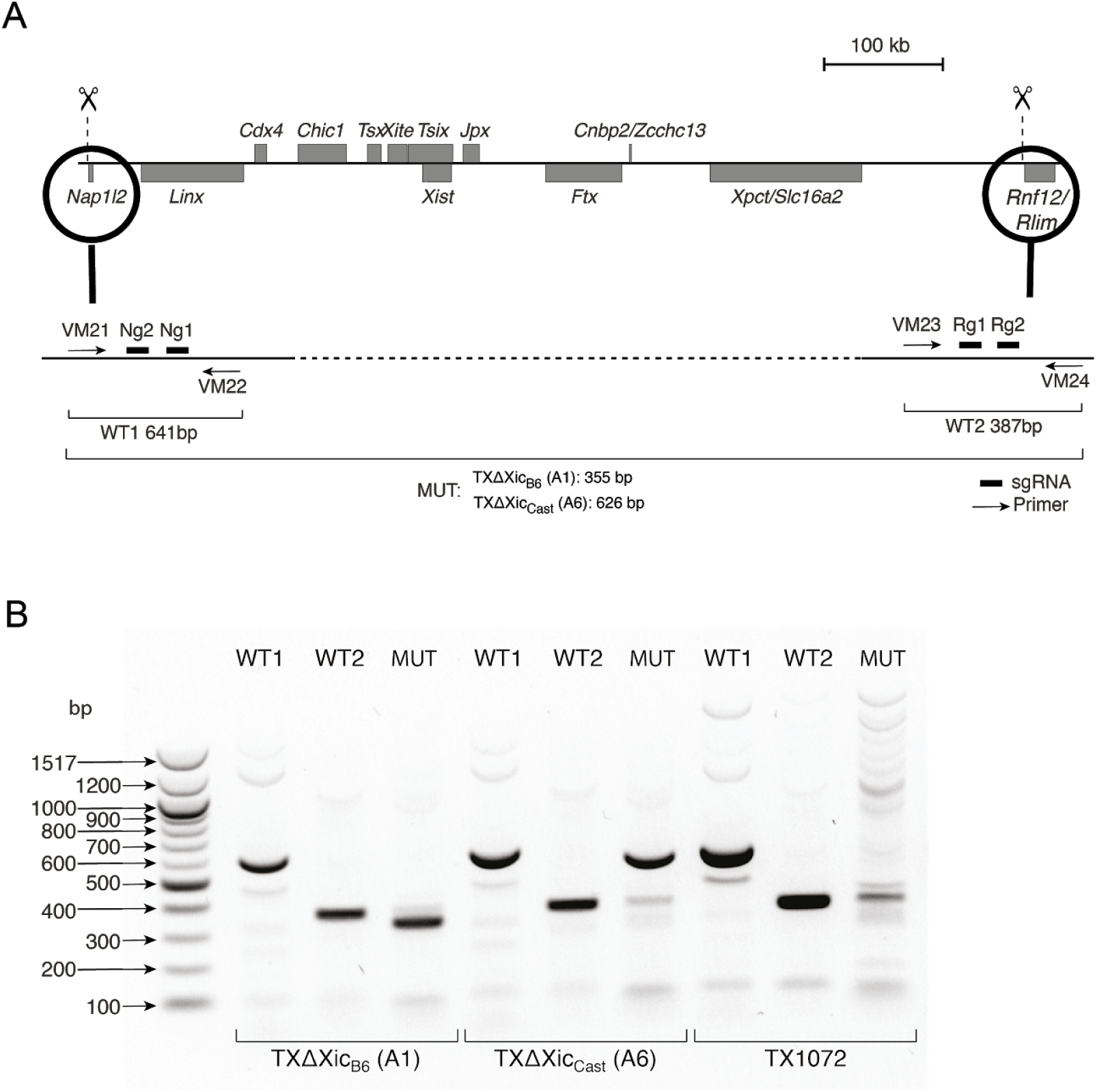
Generation of Xic deletion mESC lines. **(A)** Schematic representation of the X inactivation center (top). Genes on the plus strand are shown above the line and genes on the minus strand below. Scissors demark the deleted region. Below the position of the sgRNA used to generate the deletion (bars) and of the primers (arrows) used for genotyping are shown together with the expected sizes for the PCR products. **(B)** Genotyping PCRs for clones A1 and A6 and the parental cell line TX1072.

**Supplemental Figure S6:**
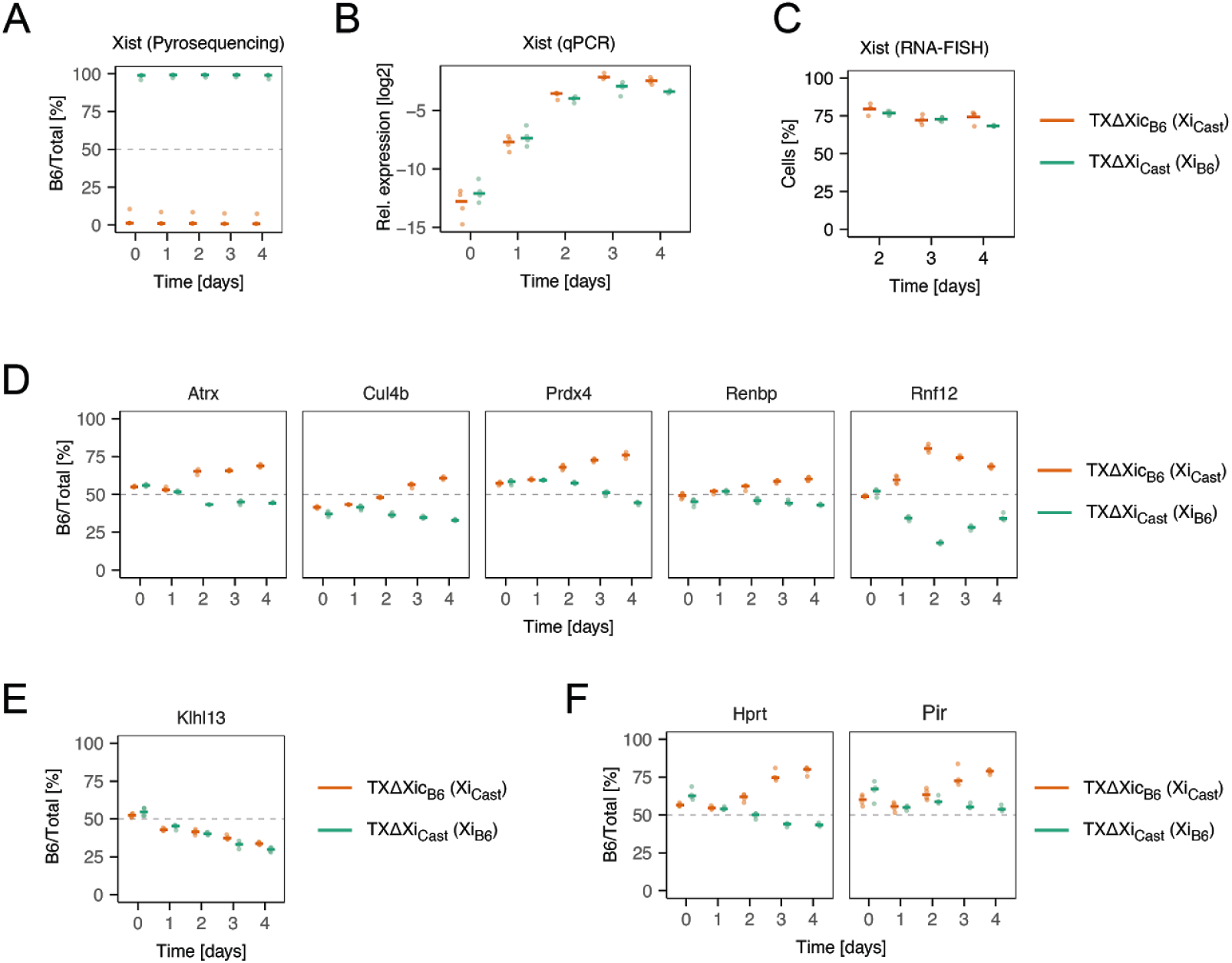
Independent validation of differential silencing dynamics. **(A-C)** Comparison of Xist expression patterns in differentiating TXΔXic_B6_ and TXΔXic_Cast_ mESCs. Allelic expression of Xist was assessed by pyrosequencing (A), relative expression by qPCR (B) and the frequency of Xist upregulation by RNA-FISH (C) for the experiments shown in Figure 6 in the main text. **(D-F)** Allelic expression of Xist assessed by pyrosequencing for all genes shown in Figure 6B-D in the main text. Bars represent the mean of 4 biological replicates, dots the individual measurements. In (C) 100 cells were analyzed for each replicate. The dashed lines A and D-F indicate the expected allelic ratio in the absence of silencing and allelic skewing.

## Supplemental Tables

**Supplemental Table S1: Cell and gene filtering**

Table summarizing the cell (Cell Filtering) and gene (Gene Filtering) filtering steps, as described in the Methods section. For both pre-processing steps, the table on top summarizes the number of cells or genes removed from the analysis for each filtering criterion.

**Supplemental Table S2: Cell and gene classification**

Classification of cells according to Xist expression, gene silencing and pseudotime and classification of X-linked genes according to their silencing behavior during XCI.

**Supplemental Table S3: Identification of putative Xist regulators**

Table summarizing the results of the differential expression analysis (MAST: Xist High vs Low) and the Spearman’s correlation analysis with Xist (Spearman: Gene CPM vs Xist CPM), aiming to identify putative Xist regulators for each time point throughout cellular differentiation. In both analyses, the FDR column represents the Benjamini-Hochberg adjusted significance values.

**Supplemental Table S4: Reagents used for generation and analysis of TX**Δ**Xic cell lines (related to Fig. 6)**

(A) Primer and sgRNA sequences for cell line generation, (B) RNA FISH probes, (C) Pyrosequencing assays, (D) qPCR primers.

